# Music-listening regulates human microRNA transcriptome

**DOI:** 10.1101/599217

**Authors:** Preethy S. Nair, Pirre Raijas, Minna Ahvenainen, Anju K. Philips, Liisa Ukkola-Vuoti, Irma Järvelä

**Author notes:** Corresponding author: Irma Järvelä.

## Abstract

Here, we used microRNA sequencing to study the effect of 20 minutes of classical music-listening on the peripheral blood microRNA transcriptome in subjects characterized for musical aptitude and music education and compared it to a control study without music for the same duration. In participants with high musical aptitude, we identified up-regulation of six microRNAs (hsa-miR-132-3p, hsa-miR-361-5p, hsa-miR-421, hsa-miR-23a-3p, hsa-miR-23b-3p, hsa-miR-25-3p) and down-regulation of two microRNAs (hsa-miR-378a-3p, hsa-miR-16-2-3p) post music-listening. The up-regulated microRNAs were found to be regulators of neuron apoptosis and neurotoxicity, consistent with previously reported neuroprotective role of music. Some up-regulated microRNAs were reported to be responsive to neuronal activity (miR-132, miR-23a, miR-23b) and modulators of neuronal plasticity, CNS myelination and cognitive functions like long-term potentiation and memory. miR-132 and *DICER*, up-regulated after music-listening, protect dopaminergic neurons and is important for retaining striatal dopamine levels. miR-23 putatively activates pro-survival PI3K/AKT signaling cascade, which is coupled with dopaminergic signaling. Some of the transcriptional regulators (FOS, CREB1, JUN, EGR1 and BDNF) of the up-regulated microRNAs are sensory-motor stimuli induced immediate early genes and top candidates associated with musical traits. Amongst these, *BDNF* is co-expressed with *SNCA*, up-regulated in music-listening and music-performance, and both are activated by GATA2, which is associated with musical aptitude. Some of the candidate microRNAs and their putative regulatory interactions were previously identified to be associated with song-learning, singing and seasonal plasticity networks in songbirds and imply evolutionary conservation of the auditory perception process: miR-23a, miR-23b and miR-25 repress *PTEN* and indirectly activates the MAPK signaling pathway, a regulator of neuronal plasticity which is activated after song-listening. We did not detect any significant changes in microRNA expressions associated with music education or low musical aptitude. Our data thereby show the importance of inherent musical aptitude for music appreciation and for eliciting the human microRNA response to music-listening.

## Introduction

Music-listening involves sensory processing of acoustic stimuli by auditory system followed by cognitive perception by the neural processing centers in the brain [1,2]. Studies based on functional brain imaging have long indicated the role of music-listening in activating the reward system and various parts of the human brain involved in the processing of emotions, learning and memory [3,4]. Music interventions have been reported to increase ergogenic effects [5], motivation [6], and have been used for motor and cognitive rehabilitation in neurological diseases [7].

Using genomics and bioinformatic approaches, we and others have shown that human musical aptitude is associated with genetic loci known to contain genes responsible for inner ear development and neurocognitive processes [8,9]. From a genetic perspective, music is an epigenetic modulator that may affect human genes and their regulation: music-listening and music-performance activated key genes expressed in songbird brain which are involved in dopaminergic neurotransmission, long-term potentiation, synaptic plasticity and memory [10,11].

Studies on Zebra finches have given strong indications that song-listening regulates both novel and known microRNAs with implications on neurogenesis and neuron differentiation [12,13]. The regulatory role played by microRNAs are well studied in nervous system development, synaptic plasticity [14,15] and amygdala-dependent memory formation [16]. MicroRNAs are also involved in inner ear development and sensory functions of the ear, hence any expression changes in the microRNA may impair the ability to hear or listen to music [17,18]. Hence, studying microRNA transcriptome changes during music-listening can give insights into post-transcriptional regulatory mechanisms associated with human sound perception and cognition. Here, we analyzed microRNA transcriptome using high-throughput sequencing and bioinformatic methods to elucidate the effect of music listening on individuals characterized for music education and musical aptitude. We used peripheral blood because brain tissues were not accessible.

## Materials and Methods

### Ethics statement

The study was approved by the ethical committee of Helsinki University Central Hospital (permission #13/03/2013) and was conducted in accordance with the Declaration of Helsinki. In addition, a written informed consent was obtained from all the subjects.

### Study population

A total of 43 volunteers of European descent attended the music-listening study and a subset of them participated in the control study (N=7). Majority of these subjects also participated to our genome-wide transcriptome profiling study [10]. The detailed phenotypic characteristics of the study participants are provided in Table 1. We wanted to evaluate the effect of music-listening based on two phenotypes, the musical aptitude and music education of the participants. Musical aptitude was defined by a combined music test score, abbreviated as *COMB* score, from tests for detecting auditory structuring ability, pitch and time; the latter two belonging to the Seashore tests for testing musical aptitude, as described before [19] (see S1 File). We sub-phenotyped the participants based on their music education and distribution of their COMB scores (range:0-148). In this study, the participants who had their COMB scores in the upper most or 4th quartile were classified as the *high COMB* participants and those below 2nd quartile as the *low COMB* participants. The data regarding music education of the participants were collected using a questionnaire. The participants have been allocated to four different Edu classes (class 1-4) based on the number of years of music education they have completed, as described before [10]. Participants in Edu classes 3 and 4 are mentioned in the study as *high Edu* group and those in Edu classes 1 or 2 have been categorized as *low Edu*.

**Table 1.** Phenotypic characteristics of the participants. Continuous explanatory variables, Age and COMB (combined musical score) are presented as the mean ± standard deviation with confidence interval at 95 % and categorical explanatory variables (gender and Edu classes) are provided as numbers.

| Category | Total<br>Subjects | Male | Female | Age | COMB<br>score | Edu<br>Low | Edu<br>High |
| --- | --- | --- | --- | --- | --- | --- | --- |
| Music<br>listening<br>Study | 43 | 20 | 23 | 42.23 +/-<br>17.09 ;<br>CI=5.26 | 130.4 +/-<br>17.12 ;<br>CI=5.27 | 18 | 25 |
| Control Study | 7 | 4 | 3 | 51.43 +/-<br>15.43 ;<br>CI=14.27 | 124.18 +/-<br>23.8 ;<br>CI=22.01 | 4 | 3 |
| High COMB | 11 | 4 | 7 | 34 +/-<br>12.77 ;<br>CI=8.58 | 145.95 +/-<br>2.19 ;<br>CI=1.47 | 1 | 10 |
| Low COMB | 11 | 6 | 5 | 60.09 +/-<br>15.48 ;<br>CI=10.4 | 105.39 +/-<br>7.83 ;<br>CI=5.26 | 8 | 3 |
| High Edu | 25 | 10 | 15 | 36.72 +/-<br>15.7 ;<br>CI=6.48 | 138.13 +/-<br>11.77 ;<br>CI=4.86 | 0 | 25 |
| Low Edu | 18 | 10 | 8 | 49.89 +/-<br>16.32 ;<br>CI=8.12 | 119.67 +/-<br>17.85 ;<br>CI=8.87 | 18 | 0 |

### Music exposure (Music-listening)

Before the study, we requested the participants not to actively listen to music on the day of the study. During the study, the participants listened to Wolfgang Amadeus Mozart’s Violin Concerto No.3 in G major, K.216 that lasted about 20 minutes in a lecture hall at Aalto University, Finland, simulating a live concert. We chose the music piece due to the popularity of Mozart music pieces in Western culture. To study the effect of music-listening on human microRNA expressions, we used a time window of 20 minutes, which is the approximate duration of one music piece in a traditional western classical concert. The duration of listening was chosen in accordance with previously reported findings from humans which indicate a time course as short as few milliseconds for the auditory processing and perception of sound [20–23]. In accordance with these, in Zebra finches, abrupt changes in the forebrain microRNA expression has been noted within 30 minutes of novel song-listening [13] as well as songs not heard for a while [12].

### Control study

The control study was organized at the Espoo Cultural Center and the participants were requested not to actively listen to music on the day of the control study. During the control study without music, the subjects (N=7) could discuss or have a walk (no exercise) outside based on their own choice for 20 minutes.

### Sample collection, microRNA extraction and sequencing

Peripheral whole blood samples were collected immediately before (Pre), as well as, immediately after (Post) 20 minutes of music-listening and control study. Details on microRNA extraction is provided in the S1 File. Sequencing libraries were prepared at the High-Throughput Genomics department of The Wellcome Trust Center for Human Genetics followed by sequencing with Illumina HiSeq.

### Pre-processing of microRNA sequencing reads

We assessed the quality of microRNA sequencing reads with FastQC version 11.3 (http://www.bioinformatics.babraham.ac.uk/projects/fastqc/). Next, we trimmed sequencing adapters from the 3’ end of 50 bp reads requiring adapter overlap of 5 bp, error rate of 0.1 and filtered shorter (<15 bp) and low-quality reads (Phred score <20) with Trim Galore! version 0.3.7 (http://www.bioinformatics.babraham.ac.uk/projects/trim_galore/). Trimmed reads were quality checked again using FastQC and aligned to human genome reference (GRCh38, Ensembl release 76) with bowtie version 1.1.2 [24]. Only unique alignments were opted for from the best alignments (–best –strata), requiring complete match for a seed length of 18. We next quantified microRNA expression using HTSeq version 0.6.1p1 [25] according to miRBase release 21 annotations for human microRNAs [26].

### Differential expression analyses of microRNA

To understand the effect of music-listening on microRNA expressions, we used DESeq2 (version 1.20.0) [27] and analyzed the differential expression of microRNAs over time (Post versus (vs.) Pre) in the music-listening group compared to the control group. Strengths of DESeq2 include its high sensitivity for experiments with a wide range of sample numbers (small to large) and for those with a small fold change [27,28]. Furthermore, benchmark comparison of statistical tools for analyzing differential expression shows DESeq2 giving a false positive rate as low as 0 and true positive rate above 80%, even with a log fold threshold and a replicate number as low as 0.5 and 6 respectively [29] and is hence the ideal tool for our study.

We performed generalized linear model based differential expression analyses with DESeq2, implementing likelihood ratio tests with a design matrix controlled for paired experimental design. False discovery rate (FDR) adjusted p-values were calculated using the Benjamini-Hochberg method that accounts for multiple testing correction. MicroRNAs were considered to be differentially expressed when FDR adjusted p-values were less than 10% [13,30]. We kept the fold change threshold of 1.2 in accordance with gene-environmental interaction studies where moderate changes in microRNA expressions have been observed [31,32]. Finally, only those DE microRNAs that showed a Post-Pre threshold of at least 10% for the music-listening session were chosen as candidate microRNAs for further analyses [10].

### Functional analysis of microRNAs

We performed functional enrichment analysis of the DE microRNAs using TAM 2.0 [33], a tool for microRNA specific enrichment analysis. For a given microRNA dataset, TAM 2.0 analyzes the over-representation of functional and disease annotations by comparing the input microRNAs to high quality, manually annotated reference microRNA dataset. TAM 2.0 then test whether the given microRNA dataset is over-represented or under-represented for functions, diseases, transcription factors (upstream-regulators) etc. by applying hyper-geometric test. TAM 2.0 analysis addresses the bias previously noted to be associated with the over-represented functions reported for microRNAs, when the over-representation analysis was performed solely based on target genes [34,35]. Additionally, TAM 2.0 performs comparative analysis for the up-regulated and down-regulated microRNAs together to correlate them to those dysregulated in disease conditions. We then collected the validated transcriptional regulators of the DE microRNAs from TransmiR 2.0 [36] and selected only those which were previously identified to be associated with musical traits [37], auditory perception [38–40], and those which were found to be positively selected in the human musical aptitude [41]. Next, we obtained the validated ontology annotations from miRBase, which are derived from experimentally verified miRNA:target interaction data [42]. Annotations for the closest orthologs of our DE microRNAs, as indicated by Alliance of Genome Resources as *Rattus norvegicus*, were collected from the Rat Genome Database (RGD) [43]. Furthermore, to correlate blood microRNA expression to the brain, we obtained tissue-wide expression patterns for DE microRNAs from the miRWalk2.0 [44], miRIAD [45], BBBomics [46], and literature.

### Identification of microRNA target genes

To understand the post-transcriptional gene regulatory mechanisms involving microRNAs, validated target genes that were supported by strong evidence (based on reporter assay or western blot) were obtained for the DE microRNAs from miRTarBase database (Release 7.0) [47]. We also collected predicted target genes for the DE microRNAs from TargetScan (Release 7.2) [48] and applied the below filtering criteria to reduce the false positive target genes. For the conserved and broadly conserved microRNA families, only those target genes with conserved sites having an aggregate probability of conserved targeting at least 0.2 and a total context++ score at most −0.15 were selected. For the poorly conserved DE microRNA families and those with other miRBase annotations, target genes with a total context++ score of less than −0.15 were selected [48]. Predicted target genes of the DE microRNAs with non-canonical binding were not considered for the analyses. Both the predicted and validated target genes of the DE microRNAs were then combined for further analysis and functional interpretation.

### Comparative analyses

We then compared the DE microRNAs to the song-responsive and singing-regulated microRNAs in Zebra Finch [12,13,49] for understanding the conservativeness of the microRNA regulatory mechanisms in music perception. We also compared target genes of the DE microRNAs to singing-regulated genes in song birds [39], and to the target genes of song-listening and singing responsive microRNAs [12,13,49]. Next, we calculated the significance of the overlap between the target genes of our DE microRNAs and the genes regulated by songbird singing using random sampling (without replacement) of our datasets (N=10,000) and overlap estimation for each of the re-sampled datasets. To this end, we created a dataset with behaviorally (singing) stimulated genes from songbird brain [38,39,50] and labeled it *song production cum perception gene set*. For the songbird set sampling, we used all the annotated genes from *Taeniopygia guttata* (N=17926) as Universe and sampling was performed for the same size as *song production cum perception gene set*. Human genes were sampled for the same size as the number of predicted and validated target genes of the down-regulated microRNAs using all annotated human genes as the Universe (N=20219). In the same way, we analyzed the overlap significance between the target genes of the up-regulated microRNAs from the *high COMB* group and the singing-inhibited genes from the songbird brain [39] using resampling (N=10,000).

### Integrated analysis and putative regulatory network construction

To understand the microRNA-gene regulatory mechanisms underlying music-listening in listeners with high musical aptitude, we integrated our microRNA findings with the music responsive gene expression findings from the same group [10] using IPA and the microRNA-gene interactions gathered from TargetScan, miRTarBase and literature. Only those target genes of the differentially expressed microRNAs from this study which showed inverse direction of regulation in the gene expression findings (from the same music-performance and control activity as this study) [10] were considered as microRNA-gene interactions in music-listening.

We further created a putative gene regulatory network in music-listening using Cytoscape 3.7.1 by merging our integrated results (above) with transcriptional regulatory data for microRNAs (TF-microRNA) from TransmiR 2.0, previously reported [10] statistically significant up-stream regulators of the DE genes (TF-gene), the microRNA-TF regulatory information from TargetScan/literature and findings related to song and music perception. Here, it is important to highlight the fact that a microRNA can simultaneously regulate the expression of multiple genes through their direct interactions or indirectly through the regulation of their transcriptional regulators (microRNA-TF) [51,52]. For this study, we collected the regulatory effect (activation/inhibition) of the up-stream regulator on the DE genes (TF-gene) and included only those TFs which were targeted by DE microRNA (microRNA-TF) to the putative regulatory network. From the validated TF-microRNA regulatory data from TransmiR 2.0, only those TFs which met our criteria described in the functional analysis were included to the network. This putative regulatory network was further extended with some of the functions from the microRNA enrichment analysis, literature findings and putative connecting molecules (including some validated target genes) between the microRNAs and the functions. Functional interactions between the up-regulated molecules, putatively up-regulated molecules and some of the transcriptional regulators in this network were also inferred with STRING [53].

## Results

### Read and microRNA statistics

All the reads of the music and the control studies passed FastQC quality check for A) basic statistics, B) per sequence quality scores and C) per base N content. The median number of raw reads per sample was 13,521,377.5 for the music study (range:8,577,637-19,549,067) and 9,355,225 for the control study (range: 8,423,477-11,094,252). On average, 97.75 % and 96.35 % of reads from the music-listening study and control study respectively were trimmed for sequencing adapters. Mean alignment percentage of the trimmed reads to human genome reference (Ensembl 76/GRCh38) was 83.6 % (range: 65.69 - 90.81) and 85.35 % (range: 79.85 - 87.88) respectively for the music-listening and control studies. All alignments from both the studies passed quality control. The top two most expressed microRNAs in the music-listening and control studies are hsa-miR-92a-3p and hsa-miR-451a. Further, 26 out of the 30 most expressed microRNAs in the music study samples are amongst the correspondingly most expressed microRNAs in the control samples.

### MicroRNA expression changes in the control study

We used all the control samples as a reference group, without sub-phenotype divisions, to compare music-listening responsive microRNA expressions. To facilitate this, we estimated expression differences in microRNA between *high Edu* (N=3) and *low Edu* participants (N=4) from the control study and used it as an indicator of homogeneity of the control samples. From this analysis, we did not observe any significant differences in microRNA expression between these groups thereby showing homogeneity of the control samples (data not shown).

### MicroRNA response to music-listening

At a very stringent FDR threshold of 5%, we observed statistically significant up-regulation of hsa-miR-132-3p, hsa-miR-361-5p, hsa-miR-421 and down-regulation of hsa-miR-378a-3p in the *high COMB* group after music-listening, compared to the control activity without music. Further, at a permissive significance threshold (FDR<10%), we also observed up-regulation of hsa-miR-23a-3p, hsa-miR-23b-3p, hsa-miR-25-3p and down-regulation of hsa-miR-16-2-3p in the same group. DE statistics for microRNAs that exhibited significant differential expression after music-listening in the *high COMB* group compared to the control study are given in Table S1. Genomic information for the DE microRNAs is provided in Table S2. Moreover, extensive literature search and information collected from biological databases [44,46] indicate expression of the DE microRNAs in blood brain barrier (BBB), nervous and auditory systems (Table S3). No statistically significant changes in microRNA expressions were found in *low COMB*, *high Edu* or *low Edu* groups after music-listening when compared to the control group.

### Putative functions of the DE microRNAs

Based on the analysis of the DE microRNAs using TAM 2.0 (p-value<0.05), the up-regulated microRNAs were found to be regulators of neuron apoptosis (hsa-mir-23a, hsa-mir-23b), hormone-mediated signaling pathway (hsa-mir-23a, hsa-mir-23b, hsa-mir-132), neurotoxicity (hsa-mir-25, hsa-mir-132), cell death (hsa-mir-23a, hsa-mir-25, hsa-mir-23b), wound healing (hsa-mir-23a, hsa-mir-132) and glucose metabolism (hsa-mir-23a, hsa-mir-23b) (Table S4). TAM 2.0 analysis also showed EGR1, GNRH1, USF1 and CREB1 as the top transcriptional regulators (p-value<0.05) of the up-regulated microRNAs. For the down-regulated microRNAs, angiogenesis (p-value=0.00372; hsa-mir-16-2, hsa-mir-378a), cell proliferation (p-value=0.00565; hsa-mir-16-2, hsa-mir-378a) and adiponectin signaling (p-value=0.00755; hsa-mir-378a) were the top functions from TAM 2.0 analysis (S4 Table). Comparative analysis wizard from TAM 2.0, which analyses the up-regulated and down-regulated microRNAs together, gave neuroblastoma as the top most result. The validated TF-microRNA interactions from the TransmiR 2.0 database which met our criteria that is described in the methods are provided in the Fig 1.

**Fig 1.**
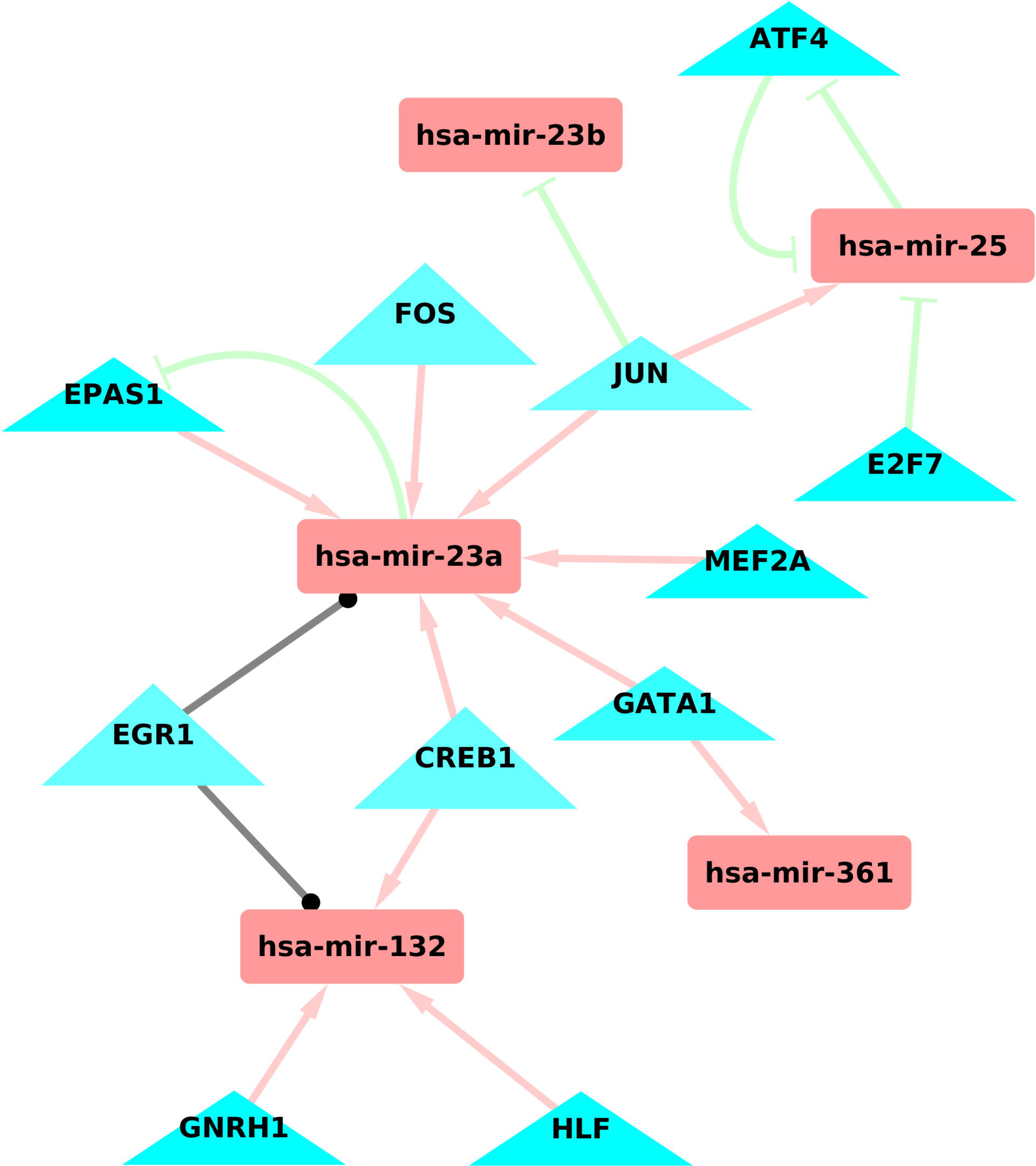
Validated transcriptional regulatory data for the DE microRNAs from TransmiR 2. In addition to the transcriptional regulators which met our criteria as per methods, we have also included GATA1 which was previously found to be associated with auditory perception.

According to the validated ontology annotations collected from miRBase, all the up-regulated microRNAs were expressed in the extracellular space (Fig 2). hsa-miR-132-2p, hsa-miR-23a-3p and hsa-miR-23b-3p share two functions: positive regulation of cell migration and endothelial cell proliferation involved in sprouting angiogenesis. hsa-miR-23a-3p and hsa-miR-23b-3p are involved in positive regulation of ERK1 and ERK2 cascade, cellular response to vascular endothelial growth factor stimulus and cell growth involved in cardiac muscle cell development. hsa-miR-132-3p was important for positive regulation of cell proliferation, vascular endothelial cell proliferation, angiogenesis, protein kinase B signaling and miRNA mediated inhibition of translation. On the contrary, hsa-miR-361-5p is involved in the negative regulation of angiogenesis; hsa-miR-25-3p is involved in the negative regulation of cardiac muscle tissue growth and stress responsive cardiac muscle hypertrophy.

**Fig 2.**
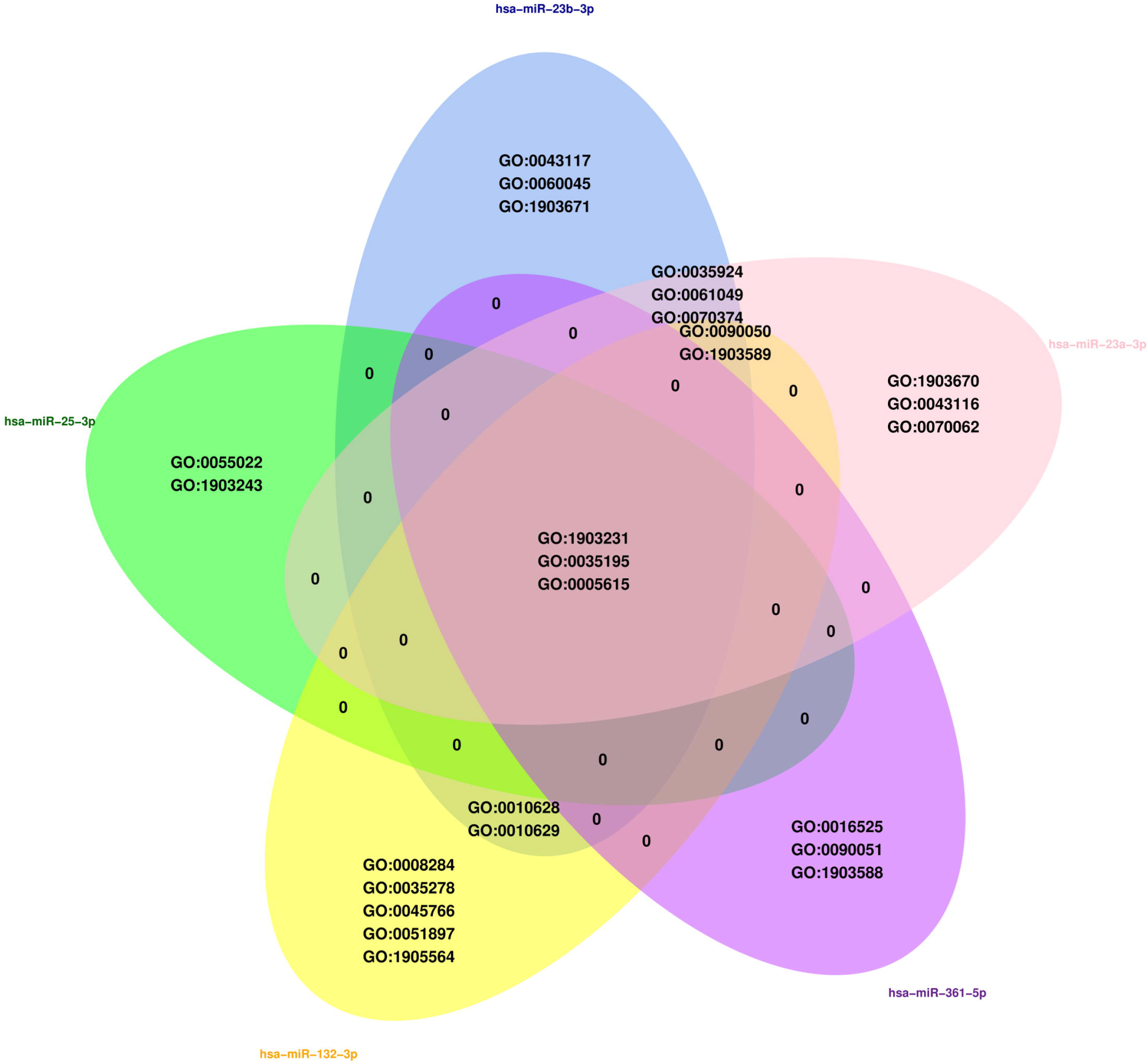
Validated ontology annotations for the DE microRNAs from miRBase. The up-regulated microRNAs, miR-23a and miR-23b play integral roles in myelination and oligodendrocyte development in the central nervous system [54] by negatively regulating lamin B1 (*LMNB1*), a repressor of oligodendrocyte and myelin specific proteins. Potential role of hsa-miR-25 and hsa-miR-23b-3p in the auditory system is supported by the expression of their host genes, *MCM7* (location: 7q22.1) and *C9orf3* respectively, in the inner ear sensory epithelia [55]. Based on literature search, hippocampal miR-132 was found to be activated during neuronal activity resulting in angiogenesis, incorporation of neurons in adults and arborization of dendrites; miR-132 knock down activated pro-inflammatory and immune signaling [56]. A concise list of the putative functions of the DE microRNAs are provided in Table 2.

**Table 2.** Putative functions of the DE microRNAs and their target genes in music-listening.

| **Up-regulated microRNAs** | **Putative functions** |
| --- | --- |
| hsa-miR-23b | CNS Myelination, long-term potentiation, dopamine signaling, auditory system development |
| hsa-miR-23a | CNS Myelination, neuronal plasticity, dopamine signaling, auditory system development |
| hsa-miR-132-3p | Activity dependent neuronal plasticity, adult neurogenesis, synaptic transmission, protection of dopaminergic neurons, songbird plasticity circuits |
| hsa-miR-361-5p | Inner ear development |
| hsa-miR-421 | Inner ear development |
| hsa-miR-25-3p | Song-learning in songbirds; neurogenesis, neuroprotection, inner ear development |
| **Target genes of up-regulated microRNAs** | **Putative Functions** |
| *CASP3, PTEN* | Neuron apoptosis |
| *CDKN1A, CDKN1B* | Cell-cycle inhibition |
| *FOXP2* | Auditory perception, sensorimotor learning, vocal plasticity |
| *LMNB1* | Repression of myelin proteins |
| **Target genes of down-regulated microRNAs** | **Putative Functions** |
| *RGS2* | Dopamine secretion and signaling; Song-learning and singing in songbirds |

As per the annotations of DE microRNA homologs from rat (*Rattus norvegicus*) [57], three of the homologs (miR-25, miR-23b and miR-132) were important for long-term synaptic potentiation and two of the homologs (miR-25 and miR-23b) for sensory perception of sound. Additionally, in *Rattus norvegicus*, miR-23b was found to be important for myelination, miR-25 for the negative regulation of apoptotic process and miR-132 for neuron maturation and dendrite development, as is found based on our literature search.

### Target genes of the DE microRNAs and their functions

Our goal with target gene finding was to understand the regulatory significance of the DE microRNAs in music-listening. We collected 147 validated human microRNA:target gene interactions for the DE microRNAs from the *high COMB* group from the miRTarBase Release 7.0 [47]. Furthermore, the predicted target genes (N=2496) from TargetScan Release 7.2 [48] for these DE microRNAs were combined with the validated targets from the same group. Notably, hsa-miR-132-3p and hsa-miR-25-3p showed validated targeting of *CDKN1A* and *CDKN1B* respectively, the cell cycle inhibitors belonging to the same family implying activation of functions like cell proliferation and differentiation in connection with music-listening; as observed in song-listening in song birds [12]. Furthermore, *PTEN*, which is a promoter of apoptotic mechanisms including that in neurons, is targeted by three of the up-regulated microRNAs from this study (hsa-miR-23a-3p, hsa-miR-23b-3p and hsa-miR-25-3p [47]),suggesting neuroprotective mechanisms associated with music-listening. Interestingly, this is consistent with the results of our microRNA specific enrichment analysis which indicated neuronal apoptosis as one of the functions regulated by the up-regulated microRNAs.

### Comparative analyses with song birds

To understand evolutionary conservation of the molecular regulatory mechanisms underlying auditory perception, we compared the DE microRNAs and their target genes to those identified in song bird song-listening and singing. Amongst the DE microRNAs, hsa-miR-25-3p, which was up-regulated in the *high COMB* group after music-listening, also showed song responsive up-regulation (tgu-miR-25) in song birds. Another DE microRNA from our study, miR-132, was found to be differentially expressed across seasons in the avian song control nuclei where its target gene network regulated cell cycle inhibitors and PTEN signaling [58]. Remarkably, miR-132 also promoted neurite outgrowth and radial migration of the neurons by its repression of *FOXP2* [59]. *FOXP2* is important for human language development and vocal learning [60,61]. The down-regulated hsa-miR-378a-3p have predicted interactions with *TLK2*, one of the predicted target genes of the song-inhibited miR-2954 in song bird [12], with functions in proliferation and neuronal differentiation. hsa-miR-378a-3p and hsa-miR-16-2-3p also show predicted interactions with song stimulated genes that are found up-regulated during song-responsive down-regulation of miR-2954 in songbirds [13]: hsa-miR-378a-3p with *RBMS1* and hsa-miR-16-2-3p with *ZDHHC17, CTNNAL1, LCOR*. Hence, the results from the comparative analysis suggest some shared molecular mechanisms relevant to the auditory perception process in songbirds and humans.

Additionally, using permutation test, we identified statistically significant overlap (p-value<0; permutation test) between target genes of the down-regulated microRNAs and the *song production cum perception gene set*. Similarly, the overlap (25.31%) that we observed between target genes of the up-regulated microRNAs from the *high COMB* group and the singing inhibited genes from songbirds [39] was found to be more than that expected by chance (re-sampling p-value=0). The target genes of the DE microRNAs that were found to be overlapping with the genes behaviorally regulated in songbirds are provided in Table S5. A schematic representation of the regulatory mechanisms associated with music-listening based on the microRNA findings is in Fig 3.

**Fig 3.**
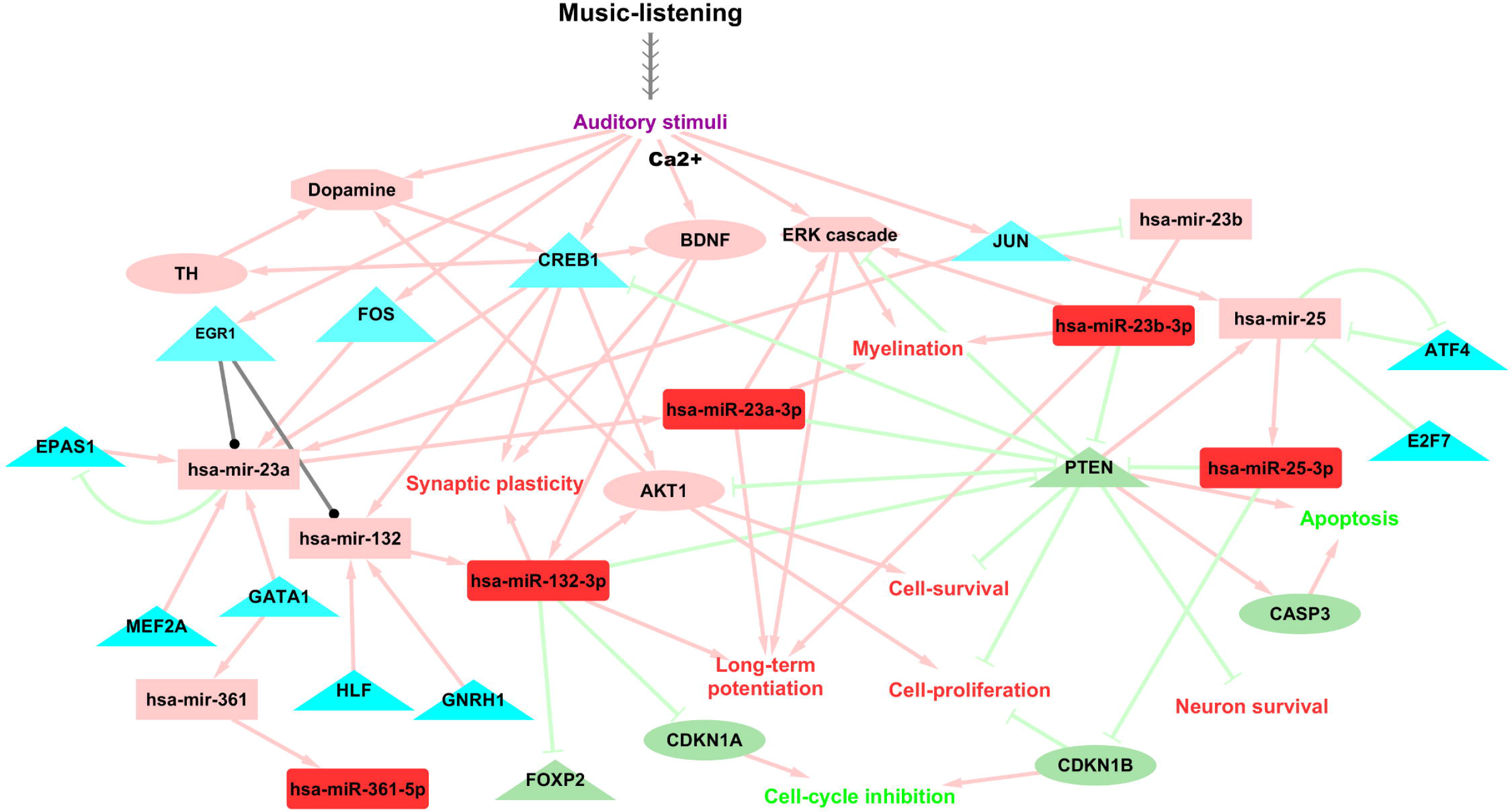
Schematic illustration of putative molecular mechanisms in music-listening based on findings from the current study. Mature microRNAs and microRNA transcripts are represented using rectangles, transcriptional regulators of the microRNAs using triangles and validated target genes using ellipses. The up-regulated molecules after music-listening are colored in red. The validated transcriptional regulators of microRNAs from TransmiR 2.0 and literature are colored in cyan. Target genes of the up-regulated microRNAs are colored in light green and includes only validated findings from the literature. Other molecules and cascades which were implied to be up-regulated based on findings from literature and from the microRNA analyses are colored in coral. Coral edges denote activation, light green edges indicate negative regulation and black lines show regulation where the direction is not known.

### Integrated results and putative regulatory network in music-listening

From the integrated analysis with gene expression data [10], we observed a total of 10 up-regulated genes in music-listening from the *high COMB* group [10] to be the target genes of 2 of the down-regulated microRNAs from the current study: hsa-miR-378a-3p shows predicted interaction with *CREBRF* and hsa-miR-16-2-3p with *UBE2B*, *SLC4A7*, *MOB1A*, *OSBPL8*, *RGS2*, *KCTD6*, *MBNL1*, *DSTN* and *TMED7*. Amongst the up-regulated microRNAs from our study, hsa-miR-132-3p was predicted to target *PSMD13*, which was found down-regulated after music-listening in the *high COMB* group [10]. Furthermore, *DICER1*, up-regulated after music-listening [10], is crucial for the biogenesis of microRNAs and functions of multiple systems [55,62,63].

A putative gene regulatory network in music-listening from the integrated analysis, extended with transcriptional regulatory data for microRNAs (TF-microRNA), TF-gene regulatory data for the DE genes, microRNA-TF interactions and findings related to song and music perception from literature is presented in Fig 4. Notably, from the merged network, we observed that hsa-miR-132-3p and hsa-miR-25-3p, which were up-regulated in the *high COMB* participants in this study, have predicted and/or functional interactions respectively with *MAPT* (Microtubule associated protein tau) and *TNFSF10* (a cytokine), two of the upstream regulators of the down-regulated genes from the same group [10]. *MAPT* is predicted to activate the down-regulated *ATP5J, HSPE1*, and *STIP1*; *MAPT* expression has been attributed to the reduced connectivity of the functional brain networks in Parkinson’s disease [64]. *TNFSF10* is predicted to activate the down-regulated *HLA-A, IFI6*, and *TNFRSF10B*. miR-132 plays a critical role in regulating TAU protein levels [65] which is important for preventing tau protein aggregation that causes Alzheimer’s disease. Furthermore, one of the down-regulated microRNAs from the *high COMB* group, hsa-miR-16-2-3p, is predicted to target *HOXA9*, one of the up-stream regulators of the genes up-regulated after music-listening in the *high COMB* group; *ADD3*, *HIGD1A*, *HNRNPU*, *MAN2A1*, *MBNL1*, and *OSBPL8*. The functional interactions between the up-regulated genes [10] and their interactants in the extended putative regulatory network (Fig 4), which were previously reported to be associated with song perception and human musical aptitude are provided in S1 Fig.

**Fig 4.**
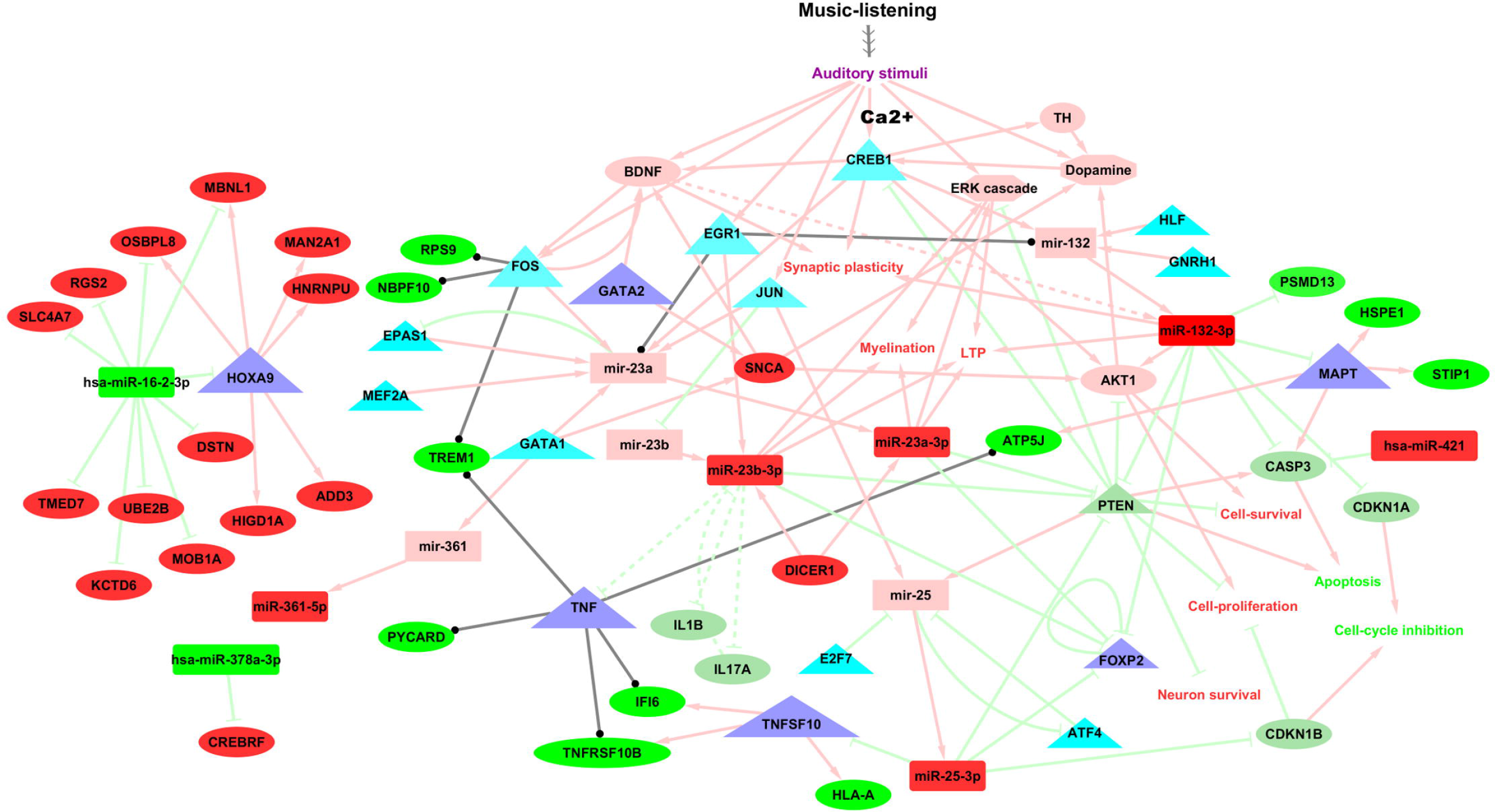
Putative gene regulatory network in music-listening. A putative gene regulatory network was constructed using Cytoscape version 3.7.1 [66] extending the results from the integrated analyses of microRNA and gene expression data after music-listening from the *high COMB* group, validated transcriptional regulators of DE microRNAs from TransmiR 2.0, validated target genes of the DE microRNAs and functions based on microRNA over-representation analysis and literature findings. Legend: Rectangles indicate microRNAs and ellipses indicate genes. The up-regulated and down-regulated molecules after music-listening are colored respectively in red and green. The validated transcriptional regulators of microRNAs from TransmiR 2.0 are colored in cyan and the upstream regulators of the DE genes from the gene expression study [10] in blue. The target genes of up-regulated microRNAs which are not differentially regulated after music-listening are colored in light green and includes only validated findings from the literature. Likewise, the target genes of down-regulated microRNAs which are not differentially regulated after music-listening are colored in coral and includes also some predicted target genes which are mentioned in the discussion. Solid lines denote direct interactions, dotted lines indicate indirect interactions, coral lines denote positive regulation,light-green lines for negative regulation and black lines show regulation where the direction is not known.

## Discussion

Here we show that musical aptitude affects peripheral blood human microRNA profiles. Perhaps the most interesting and biologically relevant results are the up-regulation of the brain-enriched microRNA, hsa-miR-132-3p, the oligodendrocyte specific microRNAs hsa-miR-23a, hsa-miR-23b and the song-stimulated miR-25 in response to music-listening in the *high COMB* group.

The up-regulation of microRNAs which are regulators of neuron apoptosis and neurotoxicity is largely consistent with early findings of neuroprotective role for music [7,67] and functions associated with musical traits [37]. For example, miR-23a and miR-23b have been experimentally confirmed to show neuroprotective effect via its repression of *APAF1*, which is an activator of caspases and neuronal apoptotic processes [33]. miR-25 and its cluster members inhibit the pro-apoptotic *TP53* and its mediators and reduce neuronal apoptotic process [68]. miR-25 also promote neurogenesis and differentiation of adult neurons by regulating TGFB-signaling pathway, that repress neurogenesis and neuronal cell proliferation, and by activating insulin-like growth factor-1 (IGF) signaling via its targeting of *PTEN*[69]. This would be consistent with our previous findings [10] which indicated that the genes down-regulated after music-listening from the *high COMB* group as activators of peptidase, endopeptidase and caspase activities. Interestingly, *PTEN* is targeted by other up-regulated microRNAs, miR-132, miR-23b and miR-23a; of these, miR-23a, through its regulation of *PTEN*, activates AKT (Protein Kinase B) signaling, PI3K (phosphatidylinositol 3-kinase) signaling, MAPK activity and promotes the expression of myelin genes [70]. MAPK signaling pathway has crucial role in the regulation of neuronal transcription, synaptic plasticity, memory consolidation [71] and was previously reported to be activated by microRNA regulation in response to song-listening in songbirds [13] implying shared pathways in auditory perception process.

Our findings also point to activity dependent regulation of the microRNAs, neuronal plasticity and augmented cognitive functions like long-term potentiation and memory after music-listening. These are in agreement with previous reports of brain plasticity and improved cognitive functions like learning, verbal ability and memory after music exposure and training [72–75]. Acoustic stimuli from music-listening induce neural spike patterns which are transduced via the auditory pathway within milliseconds [76–78] and evoke emotions in the limbic system [79]. At the molecular level, neural stimulation is conducted via calcium channel activity and neurotransmitters, which activate immediate early genes (IEG) thereby regulating gene and microRNA expression patterns [80–83]. Amongst our candidate microRNAs, miR-132 is an activity dependent microRNA which responds immediately to neuronal stimulation; miR-132 is also activated by *CREB* [84], *BDNF*, a neurotrophin which is a target of *CREB* [81], and external stimulants like cocaine. miR-132 and CREB are important for maturation and plasticity of dendrites [85]; CREB is also critical for consolidation of long-term memory and is stimulated by song-learning [86]. Interestingly, ARC, which is co-expressed with miR-132 after induction of long-term potentiation is also activated by *BDNF* [87]. It is noteworthy here that miR-23a, another candidate microRNA, is also induced by long-term potentiation with implications in memory consolidation [88]. More importantly, brain expressions of *BDNF* connected to behavioral activation of dopaminergic neurons showed positive correlation with that of *SNCA* [89,90], the candidate gene up-regulated in the *high COMB* and *high Edu* groups after music-listening [10] and in musicians after music-performance [11]. Furthermore, *SNCA* also activates *BDNF* [91] and *BDNF* and *SNCA* are regulated by *GATA2*, which is located in the strongest associated region for musical aptitude [8,10].

Of note, some of the transcriptional regulators of the candidate up-regulated microRNAs (*FOS*, *CREB1*, *JUN*, *EGR1* and *BDNF*) are activity-dependent IEGs, that are stimulated in the brain in response to auditory and motor stimuli such as songbird singing and song-listening [38,86]. Besides, *FOS*, *EGR1*, *BDNF* and *SNCA* were some of the top candidate genes associated with musical traits based on convergent evidence [37]. *BDNF* augments neurogenesis and cognition [92] and is found to be activated after music exposure [75] and songbird singing [38]. *FOS* is activated after music-performance in musicians [11] and has roles in neurotransmission and experience dependent neuroplasticity [93].

Some of the putatively activated signaling pathways and regulatory mechanisms are also indicative of dopamine signaling and protection of dopaminergic neurons. Putative activation of the pro-survival PI3K/AKT signaling cascade indirectly by music-induced miR-23a is one candidate mechanism which explains dopaminergic neurotransmission in our current study. PI3K/AKT signaling is activated in response to growth factors and neurotrophins and is coupled to the dopaminergic signaling; PI3K/AKT signaling also protects adult dopaminergic neurons from apoptosis [94]. Moreover, genetic variation in the AKT1 gene affect neural structures of the frontostriatal dopaminergic brain networks, bioavailable dopamine levels and cognitive functions [95]. *DICER*, up-regulated after music-performance [10] is another candidate molecule that protects adult dopaminergic neurons [96]. *DICER* is also critical for the maintenance of proper levels of striatal dopamine; depletion of *Dicer* in the dopamine neurons caused their progressive degeneration and loss of striatal dopamine [63]. These findings might explain the prior observations of music-listening responsive dopamine release and activation of reward pathways [4,97,98].

Our DE microRNAs and candidate molecules were also implicated in degeneration of inner ear hair cell and neurodegenerative diseases like Parkinson’s disease and Alzheimer’s disease. *DICER* is important for the biogenesis of microRNAs and functions of various systems including that of the inner ear and brain, key areas for the reception and perception of auditory signals [55,62]. Sensory neuronal *Dicer* knock out reduced the expressions of the music-induced miR-23a and miR-23b [62] and *DICER* ablation in the inner ear hair cells led to hair cell degeneration and hearing loss [55]. One of our candidate microRNAs, miR-132, protects dopaminergic neurons by its regulation of caspase3 (*CASP3*) [99], whose activated neuronal expression has been linked to dopaminergic neuronal loss of Parkinson’s disease patients [100]. A deficiency of miR-132 impaired tau metabolism which lead to tau hyperphosphorylation and aggregation, that is observed in various neurodegenerative disorders. In particular, individuals with Alzheimer’s and mild cognitive impairment had lower expression levels of miR-132 in the hipppocampal and cortical areas which declined their cognitive capabilities including working memory and perceptual speed [65]. Music listening seems to produce opposite effect by up-regulating neuroprotective microRNAs and molecules linked to neurodegenerative diseases.

Some of the candidate microRNAs and their putative regulatory interactions have previously been identified to be associated with song-learning, singing and seasonal plasticity networks in songbirds and imply conservation of the regulatory mechanisms underlying auditory perception. [12,49,58]. For instance, up-regulation of miR-25 is consistent with findings from songbirds in which it was found activated in response to song-learning and listening [12]. miR-132, which was found to be seasonally regulated in avian song control nuclei was important for sensorimotor neuronal plasticity [58]. miR-23b and miR-23a are involved in feedback regulatory circuits with its transcriptional regulator *EGR1*, an IEG that is induced in bird song learning and singing [38]. Furthermore, DE microRNAs from this study (hsa-miR-23a-3p, hsa-miR-23b-3p, hsa-miR-132- 3p, hsa-miR-25-3p) show validated and predicted targeting of *FOXP2*. Interestingly, FOXP2 is a candidate associated with music abilities including recognition and production of sound [37] and was found to be positively selected in the human evolution [41]. In songbirds, *FOXP2* is enriched in corticostriatal circuits and shows down-regulation during the sensorimotor learning period, vocal practice and after undirected singing [49,61,101]. This behavioral regulation of *FOXP2* plausibly fine tunes neural structures for learning [61] and vocal complexity [101]; suggesting here a regulatory role of the candidate microRNAs in the plasticity circuits associated with music-listening.

In this study, music-listening affected microRNA regulation only in subjects with relatively high music test scores (*high COMB*). This is in accordance with findings from the transcriptome study which identified more changes in the *high COMB* participants than in the *high Edu* group in response to the same music exposure and duration [10]. We hypothesize that drive for music is facilitated by musical aptitude; i.e., the ability to perceive the structure, pitch and duration of notes in music; and affects the human microRNA response to music-listening. Being aware of the complex cognitive trait of music, further empirical studies have to be performed in various settings, populations and using multiple genres of music to evaluate the effect of music-listening.

The high similarity (>80%) between the human brain and blood transcriptome [102] and the positive correlation of observed transcriptome changes between human cortex and whole blood [103,104] explain the rationale in our usage of peripheral blood as a representative tissue for analyzing transcriptome changes in the brain. Supporting this, a recent study on song-listening response in zebra finch shows positive correlation in the transcriptome changes of the auditory forebrain and peripheral blood [105]. Of particular interest, tissue-wide comparative analyses of microRNA expression patterns in humans indicated blood as the most similar tissue to brain [106,107]. Additionally, all the DE microRNAs in this study show expression in the blood brain barrier [46] and various parts of the nervous system (Table S3) suggestive of their cross-talk between brain and peripheral blood expressions.

## Conclusions

MicroRNAs are essential for neuronal development, plasticity and brain functions. Alterations in gene expression and regulation are important for nervous system adaptation. We hypothesize that musical aptitude serves as a pre-requisite for appreciating music and can affect the human microRNA response to music-listening. Here, the identified microRNAs provide some new insights into post-transcriptional regulatory mechanisms underlying human auditory perception and cognition. Some of the DE microRNAs shared signaling pathways with songbirds further suggesting evolutionary conservation of molecular regulatory mechanisms that regulates auditory perception. Future studies are needed to experiment with the duration of listening, genre of the music, personal preference of the participants, as well as ambiance in different combinations to understand the effect of each of these factors on microRNA expressions.

## Acknowledgements

We thank the participants for their generous collaboration. We are grateful to Petri Myllynen, Sanna Pyy, Laura Salmela, Sonja Suhonen, Jaana Oikkonen and Kai Karma for their help in organizing the study. We are grateful to Professor Harri Lähdesmäki for discussions on the study design and supervision of bioinformatic analyses. We thank Dr. Eija Korpelainen, CSC, Finland for the insightful comments that helped to improve the manuscript. We thank the High-Throughput Genomics Group at the Wellcome Trust Center for Human Genetics for the generation of the sequencing data (grant reference 090532/Z/09/Z and MRC Hub grant G0900747 91070).

## Author Contributions

P.S.N. performed data analyses, interpreted the findings, prepared figures and tables, wrote the manuscript. M.A. performed laboratory experiments and revised the manuscript. A.K.P. performed laboratory experiments. P. R. and L.U.V. contributed to the study materials. I.J. conceived the idea of the study, coordinated the study and wrote the manuscript. All authors reviewed the manuscript.

## Supporting information

**Supplemental Table S1**. Differentially expressed microRNAs from the music-listening versus control comparison in participants with high musical aptitude using analysis with DESeq2 [27]. The columns 1-3 show the DE microRNA, fold change and adjusted p-value for music-listening versus control study comparison.

**Supplemental Table S2**. Genomic location of DE microRNA transcript from HGNC and miRIAD database; conservation status, microRNA family information and number of predicted target genes of DE microRNAs from TargetScan Release 7.2; and microRNA cluster information from miRBase are given.

**Supplemental Table S3**. Candidate microRNAs associated with music-listening in participants with high musical aptitude (*high COMB*), their expression patterns and relevant functional information.

**Supplemental Table S4**. Over-represented functions for the DE microRNAs from their analysis using TAM 2.0.

**Supplemental Table S5**. Target genes of DE microRNAs that overlaps with singing and song-learning regulated genes from songbirds.

**S1 File. Supplemental Methods**.

## References

1. Menon V, Levitin DJ. The rewards of music listening: Response and physiological connectivity of the mesolimbic system. NeuroImage. 2005;28: 175–184. doi:10.1016/j.neuroimage.2005.05.053

2. Koelsch S. Brain correlates of music-evoked emotions. Nature Reviews Neuroscience. 2014;15: 170–180. doi:10.1038/nrn3666

3. Blood AJ, Zatorre RJ. Intensely pleasurable responses to music correlate with activity in brain regions implicated in reward and emotion. Proceedings of the National Academy of Sciences of the United States of America. 2001;98: 11818–11823. doi:10.1073/pnas.191355898

4. Salimpoor VN, Benovoy M, Larcher K, Dagher A, Zatorre RJ. Anatomically distinct dopamine release during anticipation and experience of peak emotion to music. Nature Neuroscience. 2011;14: 257–262. doi:10.1038/nn.2726

5. Lane AM, Davis PA, Devonport TJ. Effects of music interventions on emotional States and running performance. Journal of Sports Science & Medicine. 2011;10: 400–407.

6. Karageorghis CI, Terry PC, Lane AM, Bishop DT, Priest D-l. The BASES Expert Statement on use of music in exercise. Journal of Sports Sciences. 2012;30: 953–956. doi:10.1080/02640414.2012.676665

7. Sihvonen AJ, Särkämö T, Leo V, Tervaniemi M, Altenmüller E, Soinila S. Music-based interventions in neurological rehabilitation. The Lancet Neurology. 2017;16: 648–660. doi:10.1016/S1474-4422(17)30168-0

8. Oikkonen J, Huang Y, Onkamo P, Ukkola-Vuoti L, Raijas P, Karma K, et al. A genome-wide linkage and association study of musical aptitude identifies loci containing genes related to inner ear development and neurocognitive functions. Molecular Psychiatry. 2015;20: 275–282. doi:10.1038/mp.2014.8

9. Park H, Lee S, Kim H-J, Ju YS, Shin J-Y, Hong D, et al. Comprehensive genomic analyses associate UGT8 variants with musical ability in a Mongolian population. Journal of Medical Genetics. 2012;49: 747–752. doi:10.1136/jmedgenet-2012-101209

10. Kanduri C, Raijas P, Ahvenainen M, Philips AK, Ukkola-Vuoti L, Lähdesmäki H, et al. The effect of listening to music on human transcriptome. PeerJ. 2015;3: e830. doi:10.7717/peerj.830

11. Kanduri C, Kuusi T, Ahvenainen M, Philips AK, Lähdesmäki H, Järvelä I. The effect of music performance on the transcriptome of professional musicians. Scientific Reports. 2015;5: 9506. doi:10.1038/srep09506

12. Gunaratne PH, Lin Y-C, Benham AL, Drnevich J, Coarfa C, Tennakoon JB, et al. Song exposure regulates known and novel microRNAs in the zebra finch auditory forebrain. BMC genomics. 2011;12: 277. doi:10.1186/1471-2164-12-277

13. Lin Y-C, Balakrishnan CN, Clayton DF. Functional genomic analysis and neuroanatomical localization of miR-2954, a song-responsive sex-linked microRNA in the zebra finch. Front Neurosci. 2014;8: 409. doi:10.3389/fnins.2014.00409

14. Fineberg SK, Kosik KS, Davidson BL. MicroRNAs potentiate neural development. Neuron. 2009;64: 303–309. doi:10.1016/j.neuron.2009.10.020

15. Cohen JE, Lee PR, Chen S, Li W, Fields RD. MicroRNA regulation of homeostatic synaptic plasticity. Proceedings of the National Academy of Sciences of the United States of America. 2011;108: 11650–11655. doi:10.1073/pnas.1017576108

16. Griggs EM, Young EJ, Rumbaugh G, Miller CA. MicroRNA-182 regulates amygdala-dependent memory formation. The Journal of Neuroscience: The Official Journal of the Society for Neuroscience. 2013;33: 1734–1740. doi:10.1523/JNEUROSCI.2873-12.2013

17. Patel M, Hu BH. MicroRNAs in inner ear biology and pathogenesis. Hearing Research. 2012;287: 6–14. doi:10.1016/j.heares.2012.03.008

18. Ushakov K, Rudnicki A, Avraham KB. MicroRNAs in sensorineural diseases of the ear. Frontiers in Molecular Neuroscience. 2013;6: 52. doi:10.3389/fnmol.2013.00052

19. Pulli K, Karma K, Norio R, Sistonen P, Göring HHH, Järvelä I. Genome-wide linkage scan for loci of musical aptitude in Finnish families: Evidence for a major locus at 4q22. Journal of Medical Genetics. 2008;45: 451–456. doi:10.1136/jmg.2007.056366

20. Geiser E, Ziegler E, Jancke L, Meyer M. Early electrophysiological correlates of meter and rhythm processing in music perception. Cortex; a Journal Devoted to the Study of the Nervous System and Behavior. 2009;45: 93–102. doi:10.1016/j.cortex.2007.09.010

21. Pell MD, Kotz SA. On the time course of vocal emotion recognition. PloS One. 2011;6: e27256. doi:10.1371/journal.pone.0027256

22. Picton T. Hearing in time: Evoked potential studies of temporal processing. Ear and Hearing. 2013;34: 385–401. doi:10.1097/AUD.0b013e31827ada02

23. Rigoulot S, Pell MD, Armony JL. Time course of the influence of musical expertise on the processing of vocal and musical sounds. Neuroscience. 2015;290: 175–184. doi:10.1016/j.neuroscience.2015.01.033

24. Langmead B, Trapnell C, Pop M, Salzberg SL. Ultrafast and memory-efficient alignment of short DNA sequences to the human genome. Genome Biology. 2009;10: R25. doi:10.1186/gb-2009-10-3-r25

25. Anders S, Pyl PT, Huber W. HTSeq–a Python framework to work with high-throughput sequencing data. Bioinformatics (Oxford, England). 2015;31: 166–169. doi:10.1093/bioinformatics/btu638

26. Kozomara A, Griffiths-Jones S. miRBase: Annotating high confidence microRNAs using deep sequencing data. Nucleic Acids Research. 2014;42: D68–73. doi:10.1093/nar/gkt1181

27. Love MI, Huber W, Anders S. Moderated estimation of fold change and dispersion for RNA-seq data with DESeq2. Genome Biology. 2014;15: 550. doi:10.1186/s13059-014-0550-8

28. Zhou X, Lindsay H, Robinson MD. Robustly detecting differential expression in RNA sequencing data using observation weights. Nucleic Acids Research. 2014;42: e91. doi:10.1093/nar/gku310

29. Schurch NJ, Schofield P, Gierliński M, Cole C, Sherstnev A, Singh V, et al. How many biological replicates are needed in an RNA-seq experiment and which differential expression tool should you use? RNA (New York, NY). 2016;22: 839–851. doi:10.1261/rna.053959.115

30. Juhila J, Sipilä T, Icay K, Nicorici D, Ellonen P, Kallio A, et al. MicroRNA expression profiling reveals miRNA families regulating specific biological pathways in mouse frontal cortex and hippocampus. PloS One. 2011;6: e21495. doi:10.1371/journal.pone.0021495

31. Chilton WL, Marques FZ, West J, Kannourakis G, Berzins SP, O’Brien BJ, et al. Acute exercise leads to regulation of telomere-associated genes and microRNA expression in immune cells. PloS One. 2014;9: e92088. doi:10.1371/journal.pone.0092088

32. Tonevitsky AG, Maltseva DV, Abbasi A, Samatov TR, Sakharov DA, Shkurnikov MU, et al. Dynamically regulated miRNA-mRNA networks revealed by exercise. BMC physiology. 2013;13: 9. doi:10.1186/1472-6793-13-9

33. Li J, Han X, Wan Y, Zhang S, Zhao Y, Fan R, et al. TAM 2.0: Tool for MicroRNA set analysis. Nucleic Acids Research. 2018;46: W180–W185. doi:10.1093/nar/gky509

34. Bleazard T, Lamb JA, Griffiths-Jones S. Bias in microRNA functional enrichment analysis. Bioinformatics (Oxford, England). 2015;31: 1592–1598. doi:10.1093/bioinformatics/btv023

35. Godard P, Eyll J van. Pathway analysis from lists of microRNAs: Common pitfalls and alternative strategy. Nucleic Acids Research. 2015;43: 3490–3497. doi:10.1093/nar/gkv249

36. Tong Z, Cui Q, Wang J, Zhou Y. TransmiR v2.0: An updated transcription factor-microRNA regulation database. Nucleic Acids Research. 2019;47: D253–D258. doi:10.1093/nar/gky1023

37. Oikkonen J, Onkamo P, Järvelä I, Kanduri C. Convergent evidence for the molecular basis of musical traits. Scientific Reports. 2016;6: 39707. doi:10.1038/srep39707

38. Wada K, Howard JT, McConnell P, Whitney O, Lints T, Rivas MV, et al. A molecular neuroethological approach for identifying and characterizing a cascade of behaviorally regulated genes. Proceedings of the National Academy of Sciences of the United States of America. 2006;103: 15212–15217. doi:10.1073/pnas.0607098103

39. Whitney O, Pfenning AR, Howard JT, Blatti CA, Liu F, Ward JM, et al. Core and region-enriched networks of behaviorally regulated genes and the singing genome. Science (New York, NY). 2014;346: 1256780. doi:10.1126/science.1256780

40. Pfenning AR, Hara E, Whitney O, Rivas MV, Wang R, Roulhac PL, et al. Convergent transcriptional specializations in the brains of humans and song-learning birds. Science (New York, NY). 2014;346: 1256846. doi:10.1126/science.1256846

41. Liu X, Kanduri C, Oikkonen J, Karma K, Raijas P, Ukkola-Vuoti L, et al. Detecting signatures of positive selection associated with musical aptitude in the human genome. Scientific Reports. 2016;6: 21198. doi:10.1038/srep21198

42. Griffiths-Jones S, Grocock RJ, Dongen S van, Bateman A, Enright AJ. miRBase: microRNA sequences, targets and gene nomenclature. Nucleic Acids Research. 2006;34: D140–144. doi:10.1093/nar/gkj112

43. Shimoyama M, De Pons J, Hayman GT, Laulederkind SJF, Liu W, Nigam R, et al. The Rat Genome Database 2015: Genomic, phenotypic and environmental variations and disease. Nucleic Acids Research. 2015;43: D743–750. doi:10.1093/nar/gku1026

44. Dweep H, Gretz N. miRWalk2.0: A comprehensive atlas of microRNA-target interactions. Nature Methods. 2015;12: 697. doi:10.1038/nmeth.3485

45. Hinske LC, França GS, Torres HAM, Ohara DT, Lopes-Ramos CM, Heyn J, et al. miRIAD-integrating microRNA inter- and intragenic data. Database: The Journal of Biological Databases and Curation. 2014;2014. doi:10.1093/database/bau099

46. Kalari KR, Thompson KJ, Nair AA, Tang X, Bockol MA, Jhawar N, et al. BBBomics-Human Blood Brain Barrier Transcriptomics Hub. Frontiers in Neuroscience. 2016;10: 71. doi:10.3389/fnins.2016.00071

47. Chou C-H, Shrestha S, Yang C-D, Chang N-W, Lin Y-L, Liao K-W, et al. miRTarBase update 2018: A resource for experimentally validated microRNA-target interactions. Nucleic Acids Research. 2018;46: D296–D302. doi:10.1093/nar/gkx1067

48. Agarwal V, Bell GW, Nam J-W, Bartel DP. Predicting effective microRNA target sites in mammalian mRNAs. eLife. 2015;4. doi:10.7554/eLife.05005

49. Shi Z, Luo G, Fu L, Fang Z, Wang X, Li X. miR-9 and miR-140-5p target FoxP2 and are regulated as a function of the social context of singing behavior in zebra finches. The Journal of Neuroscience: The Official Journal of the Society for Neuroscience. 2013;33: 16510–16521. doi:10.1523/JNEUROSCI.0838-13.2013

50. Hilliard AT, Miller JE, Fraley ER, Horvath S, White SA. Molecular microcircuitry underlies functional specification in a basal ganglia circuit dedicated to vocal learning. Neuron. 2012;73: 537–552. doi:10.1016/j.neuron.2012.01.005

51. Otaegi G, Pollock A, Hong J, Sun T. MicroRNA miR-9 modifies motor neuron columns by a tuning regulation of FoxP1 levels in developing spinal cords. The Journal of Neuroscience: The Official Journal of the Society for Neuroscience. 2011;31: 809–818. doi:10.1523/JNEUROSCI.4330-10.2011

52. Li C, Zhang K, Chen J, Chen L, Wang R, Chu X. MicroRNAs as regulators and mediators of forkhead box transcription factors function in human cancers. Oncotarget. 2017;8: 12433–12450. doi:10.18632/oncotarget.14015

53. Szklarczyk D, Morris JH, Cook H, Kuhn M, Wyder S, Simonovic M, et al. The STRING database in 2017: Quality-controlled protein-protein association networks, made broadly accessible. Nucleic Acids Research. 2017;45: D362–D368. doi:10.1093/nar/gkw937

54. Lin S-T, Fu Y-H. miR-23 regulation of lamin B1 is crucial for oligodendrocyte development and myelination. Disease Models & Mechanisms. 2009;2: 178–188. doi:10.1242/dmm.001065

55. Friedman LM, Dror AA, Mor E, Tenne T, Toren G, Satoh T, et al. MicroRNAs are essential for development and function of inner ear hair cells in vertebrates. Proceedings of the National Academy of Sciences of the United States of America. 2009;106: 7915–7920. doi:10.1073/pnas.0812446106

56. Luikart BW, Bensen AL, Washburn EK, Perederiy JV, Su KG, Li Y, et al. miR-132 mediates the integration of newborn neurons into the adult dentate gyrus. PloS One. 2011;6: e19077. doi:10.1371/journal.pone.0019077

57. Smith CL, Blake JA, Kadin JA, Richardson JE, Bult CJ, Mouse Genome Database Group. Mouse Genome Database (MGD)-2018: Knowledgebase for the laboratory mouse. Nucleic Acids Research. 2018;46: D836–D842. doi:10.1093/nar/gkx1006

58. Larson TA, Lent KL, Bammler TK, MacDonald JW. William E. Wood, Caras ML, et al. Network analysis of microRNA and mRNA seasonal dynamics in a highly plastic sensorimotor neural circuit. BMC genomics. 2015;16: 905. doi:10.1186/s12864-015-2175-z

59. Clovis YM, Enard W, Marinaro F, Huttner WB, De Pietri Tonelli D. Convergent repression of Foxp2 3’UTR by miR-9 and miR-132 in embryonic mouse neocortex: Implications for radial migration of neurons. Development (Cambridge, England). 2012;139: 3332–3342. doi:10.1242/dev.078063

60. Scharff C, White SA. Genetic components of vocal learning. Annals of the New York Academy of Sciences. 2004;1016: 325–347. doi:10.1196/annals.1298.032

61. Teramitsu I, Poopatanapong A, Torrisi S, White SA. Striatal FoxP2 is actively regulated during songbird sensorimotor learning. PLoS ONE. 2010;5: e8548. doi:10.1371/journal.pone.0008548

62. Zhao J, Lee M-C, Momin A, Cendan C-M, Shepherd ST, Baker MD, et al. Small RNAs control sodium channel, expression, nociceptor excitability, and pain thresholds. The Journal of Neuroscience: The Official Journal of the Society for Neuroscience. 2010;30: 10860–10871. doi:10.1523/JNEUROSCI.1980-10.2010

63. Pang X, Hogan EM, Casserly A, Gao G, Gardner PD, Tapper AR. Dicer expression is essential for adult midbrain dopaminergic neuron maintenance and survival. Molecular and Cellular Neurosciences. 2014;58: 22–28. doi:10.1016/j.mcn.2013.10.009

64. Rittman T, Rubinov M, Vértes PE, Patel AX, Ginestet CE, Ghosh BCP, et al. Regional expression of the MAPT gene is associated with loss of hubs in brain networks and cognitive impairment in Parkinson disease and progressive supranuclear palsy. Neurobiology of Aging. 2016;48: 153–160. doi:10.1016/j.neurobiolaging.2016.09.001

65. Smith PY, Hernandez-Rapp J, Jolivette F, Lecours C, Bisht K, Goupil C, et al. miR-132/212 deficiency impairs tau metabolism and promotes pathological aggregation in vivo. Human Molecular Genetics. 2015;24: 6721–6735. doi:10.1093/hmg/ddv377

66. Shannon P, Markiel A, Ozier O, Baliga NS, Wang JT, Ramage D, et al. Cytoscape: A software environment for integrated models of biomolecular interaction networks. Genome Research. 2003;13: 2498–2504. doi:10.1101/gr.1239303

67. Conrad C. Music for healing: From magic to medicine. Lancet (London, England). 2010;376: 1980–1981.

68. Kumar M, Lu Z, Takwi AaL, Chen W, Callander NS, Ramos KS, et al. Negative regulation of the tumor suppressor p53 gene by microRNAs. Oncogene. 2011;30: 843–853. doi:10.1038/onc.2010.457

69. Brett JO, Renault VM, Rafalski VA, Webb AE, Brunet A. The microRNA cluster miR-106b~25 regulates adult neural stem/progenitor cell proliferation and neuronal differentiation. Aging. 2011;3: 108–124. doi:10.18632/aging.100285

70. Lin S-T, Huang Y, Zhang L, Heng MY, Ptácek LJ, Fu Y-H. MicroRNA-23a promotes myelination in the central nervous system. Proceedings of the National Academy of Sciences of the United States of America. 2013;110: 17468–17473. doi:10.1073/pnas.1317182110

71. Thomas GM, Huganir RL. MAPK cascade signalling and synaptic plasticity. Nature Reviews Neuroscience. 2004;5: 173–183. doi:10.1038/nrn1346

72. Roden I, Kreutz G, Bongard S. Effects of a school-based instrumental music program on verbal and visual memory in primary school children: A longitudinal study. Frontiers in Psychology. 2012;3: 572. doi:10.3389/fpsyg.2012.00572

73. Rodrigues AC, Loureiro MA, Caramelli P. Long-term musical training may improve different forms of visual attention ability. Brain and Cognition. 2013;82: 229–235. doi:10.1016/j.bandc.2013.04.009

74. Schlaug G. Musicians and music making as a model for the study of brain plasticity. Progress in Brain Research. 2015;217: 37–55. doi:10.1016/bs.pbr.2014.11.020

75. Xing Y, Chen W, Wang Y, Jing W, Gao S, Guo D, et al. Music exposure improves spatial cognition by enhancing the BDNF level of dorsal hippocampal subregions in the developing rats. Brain Research Bulletin. 2016;121: 131–137. doi:10.1016/j.brainresbull.2016.01.009

76. Bigand E, Filipic S, Lalitte P. The time course of emotional responses to music. Annals of the New York Academy of Sciences. 2005;1060: 429–437. doi:10.1196/annals.1360.036

77. Kayser C, Logothetis NK, Panzeri S. Millisecond encoding precision of auditory cortex neurons. Proceedings of the National Academy of Sciences of the United States of America. 2010;107: 16976–16981. doi:10.1073/pnas.1012656107

78. Pannese A, Grandjean D, Frühholz S. Amygdala and auditory cortex exhibit distinct sensitivity to relevant acoustic features of auditory emotions. Cortex; a Journal Devoted to the Study of the Nervous System and Behavior. 2016;85: 116–125. doi:10.1016/j.cortex.2016.10.013

79. Frühholz S, Trost W, Grandjean D. The role of the medial temporal limbic system in processing emotions in voice and music. Progress in Neurobiology. 2014;123: 1–17. doi:10.1016/j.pneurobio.2014.09.003

80. Cullinan WE, Herman JP, Battaglia DF, Akil H, Watson SJ. Pattern and time course of immediate early gene expression in rat brain following acute stress. Neuroscience. 1995;64: 477–505.

81. Tao X, Finkbeiner S, Arnold DB, Shaywitz AJ, Greenberg ME. Ca2+ influx regulates BDNF transcription by a CREB family transcription factor-dependent mechanism. Neuron. 1998;20: 709–726.

82. Neher E, Sakaba T. Multiple roles of calcium ions in the regulation of neurotransmitter release. Neuron. 2008;59: 861–872. doi:10.1016/j.neuron.2008.08.019

83. Peter M, Scheuch H, Burkard TR, Tinter J, Wernle T, Rumpel S. Induction of immediate early genes in the mouse auditory cortex after auditory cued fear conditioning to complex sounds. Genes, Brain, and Behavior. 2012;11: 314–324. doi:10.1111/j.1601-183X.2011.00761.x

84. Nudelman AS, DiRocco DP, Lambert TJ, Garelick MG, Le J, Nathanson NM, et al. Neuronal activity rapidly induces transcription of the CREB-regulated microRNA-132, in vivo. Hippocampus. 2010;20: 492–498. doi:10.1002/hipo.20646

85. Magill ST, Cambronne XA, Luikart BW, Lioy DT, Leighton BH, Westbrook GL, et al. microRNA-132 regulates dendritic growth and arborization of newborn neurons in the adult hippocampus. Proceedings of the National Academy of Sciences of the United States of America. 2010;107: 20382–20387. doi:10.1073/pnas.1015691107

86. Sakaguchi H, Wada K, Maekawa M, Watsuji T, Hagiwara M. Song-induced phosphorylation of cAMP response element-binding protein in the songbird brain. The Journal of Neuroscience: The Official Journal of the Society for Neuroscience. 1999;19: 3973–3981.

87. Wibrand K, Pai B, Siripornmongcolchai T, Bittins M, Berentsen B, Ofte ML, et al. MicroRNA regulation of the synaptic plasticity-related gene Arc. PloS One. 2012;7: e41688. doi:10.1371/journal.pone.0041688

88. Ryan B, Logan BJ, Abraham WC, Williams JM. MicroRNAs, miR-23a-3p and miR-151-3p, Are Regulated in Dentate Gyrus Neuropil following Induction of Long-Term Potentiation In Vivo. PloS One. 2017;12: e0170407. doi:10.1371/journal.pone.0170407

89. Aid-Pavlidis T, Pavlidis P, Timmusk T Meta-coexpression conservation analysis of microarray data: A “subset” approach provides insight into brain-derived neurotrophic factor regulation. BMC genomics. 2009;10: 420. doi:10.1186/1471-2164-10-420

90. Kudryavtseva NN, Bondar NP, Boyarskikh UA, Filipenko ML. Snca and Bdnf gene expression in the VTA and raphe nuclei of midbrain in chronically victorious and defeated male mice. PloS One. 2010;5: e14089. doi:10.1371/journal.pone.0014089

91. Kohno R, Sawada H, Kawamoto Y, Uemura K, Shibasaki H, Shimohama S. BDNF is induced by wild-type alpha-synuclein but not by the two mutants, A30P or A53T, in glioma cell line. Biochemical and Biophysical Research Communications. 2004;318: 113–118. doi:10.1016/j.bbrc.2004.04.012

92. Strait DL, Kraus N. Biological impact of auditory expertise across the life span: Musicians as a model of auditory learning. Hearing Research. 2014;308: 109–121. doi:10.1016/j.heares.2013.08.004

93. Kaczmarek L, Nikołajew E. C-fos protooncogene expression and neuronal plasticity. Acta Neurobiologiae Experimentalis. 1990;50: 173–179.

94. Nair VD, Olanow CW, Sealfon SC. Activation of phosphoinositide 3-kinase by D2 receptor prevents apoptosis in dopaminergic cell lines. The Biochemical Journal. 2003;373: 25–32. doi:10.1042/BJ20030017

95. Tan H-Y, Nicodemus KK, Chen Q, Li Z, Brooke JK, Honea R, et al. Genetic variation in AKT1 is linked to dopamine-associated prefrontal cortical structure and function in humans. The Journal of Clinical Investigation. 2008;118: 2200–2208. doi:10.1172/JCI34725

96. Chmielarz P, Konovalova J, Najam SS, Alter H, Piepponen TP, Erfle H, et al. Dicer and microRNAs protect adult dopamine neurons. Cell Death & Disease. 2017;8: e2813. doi:10.1038/cddis.2017.214

97. Sutoo D, Akiyama K. Music improves dopaminergic neurotransmission: Demonstration based on the effect of music on blood pressure regulation. Brain Research. 2004;1016: 255–262. doi:10.1016/j.brainres.2004.05.018

98. Moraes MM, Rabelo PCR, Pinto VA, Pires W, Wanner SP, Szawka RE, et al. Auditory stimulation by exposure to melodic music increases dopamine and serotonin activities in rat forebrain areas linked to reward and motor control. Neuroscience Letters. 2018;673: 73–78. doi:10.1016/j.neulet.2018.02.058

99. Sun Z-Z, Lv Z-Y, Tian W-J, Yang Y. MicroRNA-132 protects hippocampal neurons against oxygen-glucose deprivation-induced apoptosis. International Journal of Immunopathology and Pharmacology. 2017;30: 253–263. doi:10.1177/0394632017715837

100. Hartmann A, Hunot S, Michel PP, Muriel MP, Vyas S, Faucheux BA, et al. Caspase-3: A vulnerability factor and final effector in apoptotic death of dopaminergic neurons in Parkinson’s disease. Proceedings of the National Academy of Sciences of the United States of America. 2000;97: 2875–2880. doi:10.1073/pnas.040556597

101. Heston JB, White SA. Behavior-linked FoxP2 regulation enables zebra finch vocal learning. The Journal of Neuroscience: The Official Journal of the Society for Neuroscience. 2015;35: 2885–2894. doi:10.1523/JNEUROSCI.3715-14.2015

102. Liew C-C, Ma J, Tang H-C, Zheng R, Dempsey AA. The peripheral blood transcriptome dynamically reflects system wide biology: A potential diagnostic tool. The Journal of Laboratory and Clinical Medicine. 2006;147: 126–132. doi:10.1016/j.lab.2005.10.005

103. Sullivan PF, Fan C, Perou CM. Evaluating the comparability of gene expression in blood and brain. American Journal of Medical Genetics Part B, Neuropsychiatric Genetics: The Official Publication of the International Society of Psychiatric Genetics. 2006;141B: 261–268. doi:10.1002/ajmg.b.30272

104. Rao P, Benito E, Fischer A. MicroRNAs as biomarkers for CNS disease. Frontiers in Molecular Neuroscience. 2013;6: 39. doi:10.3389/fnmol.2013.00039

105. Louder MIM, Hauber ME, Balakrishnan CN. Early social experience alters transcriptomic responses to species-specific song stimuli in female songbirds. Behavioural Brain Research. 2018;347: 69–76. doi:10.1016/j.bbr.2018.02.034

106. Liang Y, Ridzon D, Wong L, Chen C. Characterization of microRNA expression profiles in normal human tissues. BMC genomics. 2007;8: 166. doi:10.1186/1471-2164-8-166

107. Martins M, Rosa A, Guedes LC, Fonseca BV, Gotovac K, Violante S, et al. Convergence of miRNA expression profiling, α-synuclein interacton and GWAS in Parkinson’s disease. PloS One. 2011;6: e25443. doi:10.1371/journal.pone.0025443

