## Supplemental Methods for "Music-listening regulates human microRNA transcriptome"

### Subphenotyping of study population

Unrelated study subjects from our genome-wide association studies on musical aptitude [1] were invited by electronic mail to participate in this study. Majority of these subjects also participated to our recent genome-wide transcriptional profiling study and included only individuals of European descent [2]. All participants had been assessed individually for their musical aptitude [1,3] and their music education was ascertained based on a web-based questionnaire. Shortly, participants were assigned a combined musical score, abbreviated as *COMB* score, based on three musical aptitude tests - a) test for detecting auditory structuring ability [4] and b) tests for detecting the pitch and c)time, two of which are from the conventional Seashore tests for testing musical aptitude [5] used in music studies [2,3,6]. The three test scores were combined to form combined music score, abbreviated as *COMB* score [6].

### MicroRNA extraction and sequencing

2.5ml of peripheral blood was drawn into PAXgene Blood RNA tubes (PreAnalytiX GmbH, Hombrechtikon, Switzerland) before and after listening and control studies as per kit instructions and stored at -20°C until processing. RNA was isolated using either the PAXgene Blood miRNA Kit (PreAnalytiX GmbH, Hombrechtikon, Switzerland) or Ambion's MagMAX™ for Stabilized Blood Tubes RNA Isolation Kit (Compatible with PAXgene® Blood RNA Tubes; Applied Biosystems, USA) as per the kit manual. Purified RNA samples were measured for concentration and purity on the NanoDrop 1000 v.3.7 (Thermo Fisher Scientific, USA). Pre and post-listening samples for each participant were processed using the same kit.

Samples then underwent a concentration protocol; briefly, 0.1 volume of 3M Sodium acetate buffer solution (Sigma-Aldrich, Finland), 5ug Glycogen (Sigma-Aldrich, Finland) and 3 volumes of 100% ethanol (Altia Oyj, Finland) were added to each RNA sample. Samples were mixed well then incubated at -80°C for 30 minutes before centrifuging at >12,000 x g for 30 minutes at 4°C. Supernatant was removed and 1ml ice cold 70% ethanol was added to each sample, vortexed, then centrifuged for 10 minutes at >12,000 x g at 4°C; this wash step was repeated once. Supernatant was removed and samples were incubated for 10 minutes on ice with caps open to allow remaining ethanol to evaporate before re-suspending the RNA pellet in an appropriate amount of RNase-free water (Applied Biosystems, USA). Finally, concentration and purity of the samples were measured on the NanoDrop 1000 and sample integrity was evaluated on 2100 Bioanalyzer (Agilent Technologies, Germany).

Library preparation and sequencing was performed at High Throughput Genomics department of The Wellcome Trust Center for Human Genetics, Oxford University. To be specific, samples were again checked for concentration and purity. Subsequently, small RNA libraries were prepared using NEBNext® Small RNA Library Prep Set for Illumina (Multiplex Compatible) according to the manufacturer's recommendations.

### References


1. Ukkola-Vuoti L, Oikkonen J, Onkamo P, Karma K, Raijas P, Järvelä I. Association of the arginine vasopressin receptor 1A (AVPR1A) haplotypes with listening to music. Journal of Human Genetics. 2011;56: 324–329. doi:10.1038/jhg.2011.13

2. Kanduri C, Raijas P, Ahvenainen M, Philips AK, Ukkola-Vuoti L, Lähdesmäki H, et al. The effect of listening to music on human transcriptome. PeerJ. 2015;3: e830. doi:10.7717/peerj.830

3. Pulli K, Karma K, Norio R, Sistonen P, Göring HHH, Järvelä I. Genome-wide linkage scan for loci of musical aptitude in Finnish families: Evidence for a major locus at 4q22. Journal of Medical Genetics. 2008;45: 451–456. doi:10.1136/jmg.2007.056366

4. Karma K. Musical aptitude definition and measure validation: Ecological validity can endanger the construct validity of musical aptitude tests. Psychomusicology: A Journal of Research in Music Cognition. 2007;19: 79–90. doi:10.1037/h0094033

5. Seashore CE. The Measurement of Musical Talent. The Musical Quarterly. 1915;1: 129–148. Available: http://www.jstor.org/stable/738047

6. Oikkonen J, Järvelä I. Genomics approaches to study musical aptitude. BioEssays: News and Reviews in Molecular, Cellular and Developmental Biology. 2014;36: 1102–1108. doi:10.1002/bies.201400081
