## Supplemental Table S1 for "Music-listening regulates human microRNA transcriptome"

| DE microRNA | Fold Change | Adjusted p-value |
|---|---|---|
| hsa-miR-361-5p | 1.6531877 | 0.0339218 |
| hsa-miR-378a-3p | 0.6242885 | 0.0339218 |
| hsa-miR-132-3p | 1.6905415 | 0.0384370 |
| hsa-miR-421 | 1.5911698 | 0.0444978 |
| hsa-miR-23a-3p | 1.3411910 | 0.0718555 |
| hsa-miR-23b-3p | 1.3394284 | 0.0718555 |
| hsa-miR-16-2-3p | 0.4697190 | 0.0980891 |
| hsa-miR-25-3p | 1.4865888 | 0.0991897 |
