## Supplemental Table S2 for "Music-listening regulates human microRNA transcriptome"

***Intragenic DE microRNAs***

| DE microRNA | Precursor miRNA | Family (from Targetscan 7.2 [1]) | Family Conservation from Targetscan 7.2 | Clustered miRNAs (< 10kb from the DE microRNA) from miRBase [2] | Total no. of predicted targets from Targetscan 7.2 | Band from HGNC [3] | Chromosome | Start, from miRIAD database [4] | End | Host gene |
|---|---|---|---|---|---|---|---|---|---|---|
| hsa-miR-361-5p | hsa-mir-361 | miR-361-5p | conserved | | 295 transcripts | Xq21.2 | chrX | 85903636 | 85903707 | ***CHM*** |
| hsa-miR-378a-3p | hsa-mir-378a | miR-378-3p | conserved | | 223 transcripts | 5q32 | chr5 | 149732825 | 149732890 | ***PPARGC1B*** |
| hsa-miR-23b-3p | hsa-mir-23b | miR-23-3p | broadly conserved | hsa-mir-23b, hsa-mir-27b, hsa-mir-3074, hsa-mir-24-1 | 1332 transcripts | 9q22.32 | chr9 | 95085208 | 95085304 | ***C9orf3*** |
| hsa-miR-16-2-3p | hsa-mir-16-2 | miR-16-2-3p/195-3p | other miRBase annotation | hsa-mir-15b, hsa-mir-16-2 | 4485 transcripts | 3q25.33 | chr3 | 160404745 | 160404825 | ***SMC4*** |
| hsa-miR-25-3p | hsa-mir-25 | miR-25-3p/32-5p/92-3p/363-3p/367-3p | broadly conserved | hsa-mir-106b, hsa-mir-93, hsa-mir-25 | 1037 transcripts | 7q22.1 | chr7 | 100093560 | 100093643 | ***MCM7*** |

***Intergenic DE microRNAs***

| DE microRNA | Precursor miRNA | Family (from Targetscan 7.2 [1]) | Family Conservation from Targetscan 7.2 | Clustered miRNAs (< 10kb from the DE microRNA) from miRBase [2] | Total no. of predicted targets from Targetscan 7.2 | Band from HGNC [3] | Chromosome | Start, from miRIAD database [4] | End |
|---|---|---|---|---|---|---|---|---|---|
| hsa-miR-132-3p | hsa-mir-132 | miR-132-3p/212-3p | broadly conserved | hsa-mir-132 and hsa-mir-212 | 474 transcripts | 17p13.3 | chr17 | 2049908 | 2050008 |
| hsa-miR-421 | hsa-mir-421 | miR-421 | conserved | hsa-mir-421, hsa-mir-374c, hsa-mir-374b | 451 transcripts | Xq13.2 | chrX | 74218377 | 74218461 |
| hsa-miR-23a-3p | hsa-mir-23a | miR-23-3p | broadly conserved | hsa-mir-23a, hsa-mir-27a, hsa-mir-24-2 | 1332 transcripts | 19p13.12 | chr19 | 13836587 | 13836659 |

### References

1. Agarwal V, Bell GW, Nam J-W, Bartel DP. Predicting effective microRNA target sites in mammalian mRNAs. eLife. 2015;4. doi:10.7554/eLife.05005
2. Kozomara A, Griffiths-Jones S. miRBase: Annotating high confidence microRNAs using deep sequencing data. Nucleic Acids Research. 2014;42: D68–73. doi:10.1093/nar/gkt1181
3. Yates B, Braschi B, Gray KA, Seal RL, Tweedie S, Bruford EA. Genenames.org: The HGNC and VGNC resources in 2017. Nucleic Acids Research. 2017;45: D619–D625. doi:10.1093/nar/gkw1033
4. Hinske LC, França GS, Torres HAM, Ohara DT, Lopes-Ramos CM, Heyn J, et al. miRIAD-integrating microRNA inter- and intragenic data. Database: The Journal of Biological Databases and Curation. 2014;2014. doi:10.1093/database/bau099
