## Supplemental Table S3 for "Music-listening regulates human microRNA transcriptome"

| DE microRNA | Findings on tissue expression and functions from literature |
|---|---|
| miR-23b | miR-23b is expressed in new born mouse vestibule, cochleae [1] and is located within the introns of murine cochlear and vestibular sensory epithelial coding gene, *2010111I01Rik* [2]. Hence, miR-23b is speculated to have a role in the development and functions of the inner ear [3]. miR-23b is expressed in human oligodendrocytes [4–6] and during the development of the mammalian brain in the cortex, when it was co-expressed with another music-listening regulated microRNA, miR-132 [7,8]. miR-23b was found to be under the regulation of *Dicer* and was up-regulated during myelination in the CNS and the PNS [9]. In neurodegenerative diseases like Alzheimer's disease and mild cognitive impairment, miR-23b was found down-regulated in the exosome, frontal cortex and white matter [10–12]. In mouse model for traumatic brain injury, miR-23b-3p confers neuroprotection by attenuating proapoptic Bcl-2 (B-cell lymphoma 2) proteins (*Puma* (BCL2 binding component 3), *Noxa* (phorbol-12-myristate-13-acetate-induced protein 1), and *Bax* (BCL2-associated X protein)) and neuronal loss in the hippocampus and dentate gyrus [13]. Consistent with this, a down-regulation of miR-23b has been noted in traumatic brain injury patients. Furthermore, human miR-23b-3p is predicted to interact with one of these, *PMAIP1* (*NOXA*), and *BCL2* [14]. Increased expression of miR-23b was found to mitigate autophagy of hippocampal neurons by its regulation of *ATG1* leading to improvements in cognitive abilities like memory and spatial learning [15]. miR-23b is also a regulator of pro-inflammatory cytokines (*IL-1β* (interleukin 1 beta), *IL-17* (interleukin 17A)) and TNF-mediated signaling via its targeting of *TAB2* (TGF-beta activated kinase 1 (MAP3K7) binding protein 2) and *TAB3* (TGF-beta activated kinase 1 (MAP3K7) binding protein 3) [16]. |
| miR-23a | Expressed in the inner ear [1,3], different parts of the brain (cortex, striatum and hippocampus) and during hematopoietic lineage differentiation [17]. miR-23a is a candidate microRNA for non-syndromic hereditary hearing loss (NSHHL) in humans [2]. miR-23a is characterized to be activated following *in vivo* induction of long term potentiation [18] and has important roles in the myelination of the CNS [5]. Dysregulation of miR-23a has been noted in various neurodegenerative diseases. In mild cognitive impairment and Alzheimer's, miR-23a is down-regulated in the white matter and frontal cortex [10,12]. In multiple sclerosis, miR-23a is down-regulated in the serum and peripheral T-cells [19,20]. De-myelination and oligodendrocyte damage are some of characteristics of multiple sclerosis, where miR-23a might be implicated. Increased expression of miR-23a in experimental brain ischaemia has been shown to reduce caspase-3 levels, cerebral infarction volume and oxidative stress [21]. |
| miR-132 | miR-132 is a brain enriched microRNA [22] expressed in developing mammalian brains [7,8,23]. hsa-miR-132-3p is expressed in human neurons, brain and dendritic spines [24] and is important for the development of dendritic spine density and synaptic plasticity. miR-132 is expressed in cortical neurons [25] and sensory epithelia of cochlea and vestibule [1,26]. In mouse, the miR-132 gene is located within the intron of protein-coding gene *Smg6* which is expressed in mouse inner ear sensory epithelia [2]. miR-132 is also a regulator of hippocampal BACE1 (beta-secretase 1) protein levels [27] which is involved in abnormal processing of amyloid precursor protein causing amyloid aggregation and learning disability [28]. Furthermore, miR-132 targets *Foxp2* (forkhead box P2) [29], which is crucial for vocal learning in humans and songbirds. |
| miR-361 | Expressed in the inner ear and is located within the introns of *Chm* (choroidermia (RAB escort protein 1)), protein-coding gene expressed in mouse cochlear sensory epithelia [2]. As per miRIAD database, hsa-miR-361, although expressed in many tissues, shows the highest expression in brain [30]. miR-361 is found to be down-regulated in the white matter in the neurodegenerative disease, Alzheimer's [10]. |
| miR-421 | Located within the intron of protein-coding gene expressed in mouse inner ear sensory epithelia *B230206F22Rik* [2]. miR-421 which is up-regulated after music-listening in the high COMB participants also targets *BACE1* [31], which is a catalyst for amyloid beta peptide formation from amyloid precursor protein. |
| miR-25 | Expressed in the vestibule and cochlea of newborn mouse and is located within *Mcm7* (minichromosome maintenance complex component 7) which is expressed in the inner ear sensory epithelia [2]. miR-25 is one of the candidate microRNA for non-syndromic hereditary hearing loss (NSHHL) in humans located on 7q22.1 [2]. Interestingly, a deletion of the 7q22–33 region, where miR-25 gene is located, has been associated with autism, dyslexia and atypical cerebral asymmetry [32]. Furthermore, mutations on *SLC26A5* (Prestin), which is also located in 7q22.1 and encodes a motor protein of cochlear hair cells that is necessary for amplification of sound signals, have been associated with sensorineuronal hearing loss (OMIM 613865). |

### References


1. Weston MD, Pierce ML, Rocha-Sanchez S, Beisel KW, Soukup GA. MicroRNA gene expression in the mouse inner ear. Brain Research. 2006;1111: 95–104. doi:10.1016/j.brainres.2006.07.006

2. Friedman LM, Dror AA, Mor E, Tenne T, Toren G, Satoh T, et al. MicroRNAs are essential for development and function of inner ear hair cells in vertebrates. Proceedings of the National Academy of Sciences of the United States of America. 2009;106: 7915–7920. doi:10.1073/pnas.0812446106

3. Elkan-Miller T, Ulitsky I, Hertzano R, Rudnicki A, Dror AA, Lenz DR, et al. Integration of transcriptomics, proteomics, and microRNA analyses reveals novel microRNA regulation of targets in the mammalian inner ear. PloS One. 2011;6: e18195. doi:10.1371/journal.pone.0018195

4. Letzen BS, Liu C, Thakor NV, Gearhart JD, All AH, Kerr CL. MicroRNA expression profiling of oligodendrocyte differentiation from human embryonic stem cells. PloS One. 2010;5: e10480. doi:10.1371/journal.pone.0010480

5. Lin S-T, Huang Y, Zhang L, Heng MY, Ptácek LJ, Fu Y-H. MicroRNA-23a promotes myelination in the central nervous system. Proceedings of the National Academy of Sciences of the United States of America. 2013;110: 17468–17473. doi:10.1073/pnas.1317182110

6. Lin S-T, Fu Y-H. miR-23 regulation of lamin B1 is crucial for oligodendrocyte development and myelination. Disease Models & Mechanisms. 2009;2: 178–188. doi:10.1242/dmm.001065

7. Lagos-Quintana M, Rauhut R, Yalcin A, Meyer J, Lendeckel W, Tuschl T. Identification of tissue-specific microRNAs from mouse. Current biology: CB. 2002;12: 735–739.

8. Miska EA, Alvarez-Saavedra E, Townsend M, Yoshii A, Sestan N, Rakic P, et al. Microarray analysis of microRNA expression in the developing mammalian brain. Genome Biology. 2004;5: R68. doi:10.1186/gb-2004-5-9-r68

9. Bremer J, O'Connor T, Tiberi C, Rehrauer H, Weis J, Aguzzi A. Ablation of Dicer from murine Schwann cells increases their proliferation while blocking myelination. PloS One. 2010;5: e12450. doi:10.1371/journal.pone.0012450

10. Satoh J-i. Molecular network of microRNA targets in Alzheimer's disease brains. Experimental Neurology. 2012;235: 436–446. doi:10.1016/j.expneurol.2011.09.003

11. Lugli G, Cohen AM, Bennett DA, Shah RC, Fields CJ, Hernandez AG, et al. Plasma Exosomal miRNAs in Persons with and without Alzheimer Disease: Altered Expression and Prospects for Biomarkers. PloS One. 2015;10: e0139233. doi:10.1371/journal.pone.0139233

12. Weinberg RB, Mufson EJ, Counts SE. Evidence for a neuroprotective microRNA pathway in amnestic mild cognitive impairment. Frontiers in Neuroscience. 2015;9: 430. doi:10.3389/fnins.2015.00430

13. Sabirzhanov B, Zhao Z, Stoica BA, Loane DJ, Wu J, Borroto C, et al. Downregulation of miR-23a and miR-27a following experimental traumatic brain injury induces neuronal cell death through activation of proapoptotic Bcl-2 proteins. The Journal of Neuroscience: The Official Journal of the Society for Neuroscience. 2014;34: 10055–10071. doi:10.1523/JNEUROSCI.1260-14.2014

14. Agarwal V, Bell GW, Nam J-W, Bartel DP. Predicting effective microRNA target sites in mammalian mRNAs. eLife. 2015;4. doi:10.7554/eLife.05005

15. Sun L, Liu A, Zhang J, Ji W, Li Y, Yang X, et al. miR-23b improves cognitive impairments in traumatic brain injury by targeting ATG12-mediated neuronal autophagy. Behavioural Brain Research. 2016; doi:10.1016/j.bbr.2016.09.020

16. Zhu S, Pan W, Song X, Liu Y, Shao X, Tang Y, et al. The microRNA miR-23b suppresses IL-17-associated autoimmune inflammation by targeting TAB2, TAB3 and IKK-α. Nature Medicine. 2012;18: 1077–1086. doi:10.1038/nm.2815

17. Landgraf P, Rusu M, Sheridan R, Sewer A, Iovino N, Aravin A, et al. A mammalian microRNA expression atlas based on small RNA library sequencing. Cell. 2007;129: 1401–1414. doi:10.1016/j.cell.2007.04.040

18. Ryan B, Logan BJ, Abraham WC, Williams JM. MicroRNAs, miR-23a-3p and miR-151-3p, Are Regulated in Dentate Gyrus Neuropil following Induction of Long-Term Potentiation In Vivo. PloS One. 2017;12: e0170407. doi:10.1371/journal.pone.0170407

19. Fenoglio C, Ridolfi E, Cantoni C, De Riz M, Bonsi R, Serpente M, et al. Decreased circulating miRNA levels in patients with primary progressive multiple sclerosis. Multiple Sclerosis (Houndmills, Basingstoke, England). 2013;19: 1938–1942. doi:10.1177/1352458513485654

20. Jernås M, Malmeström C, Axelsson M, Nookaew I, Wadenvik H, Lycke J, et al. MicroRNA regulate immune pathways in T-cells in multiple sclerosis (MS). BMC immunology. 2013;14: 32. doi:10.1186/1471-2172-14-32

21. Zhao H, Tao Z, Wang R, Liu P, Yan F, Li J, et al. MicroRNA-23a-3p attenuates oxidative stress injury in a mouse model of focal cerebral ischemia-reperfusion. Brain Research. 2014;1592: 65–72. doi:10.1016/j.brainres.2014.09.055

22. Sempere LF, Freemantle S, Pitha-Rowe I, Moss E, Dmitrovsky E, Ambros V. Expression profiling of mammalian microRNAs uncovers a subset of brain-expressed microRNAs with possible roles in murine and human neuronal differentiation. Genome Biology. 2004;5: R13. doi:10.1186/gb-2004-5-3-r13

23. Mukilan M, Ragu Varman D, Sudhakar S, Rajan KE. Activity-dependent expression of miR-132 regulates immediate-early gene induction during olfactory learning in the greater short-nosed fruit bat, Cynopterus sphinx. Neurobiology of Learning and Memory. 2015;120: 41–51. doi:10.1016/j.nlm.2015.02.010

24. Dweep H, Gretz N. miRWalk2.0: A comprehensive atlas of microRNA-target interactions. Nature Methods. 2015;12: 697. doi:10.1038/nmeth.3485

25. Kim J, Krichevsky A, Grad Y, Hayes GD, Kosik KS, Church GM, et al. Identification of many microRNAs that copurify with polyribosomes in mammalian neurons. Proceedings of the National Academy of Sciences of the United States of America. 2004;101: 360–365. doi:10.1073/pnas.2333854100

26. Rudnicki A, Isakov O, Ushakov K, Shivatzki S, Weiss I, Friedman LM, et al. Next-generation sequencing of small RNAs from inner ear sensory epithelium identifies microRNAs and defines regulatory pathways. BMC Genomics. 2014;15: 484. doi:10.1186/1471-2164-15-484

27. Salta E, Sierksma A, Vanden Eynden E, De Strooper B. miR-132 loss de-represses ITPKB and aggravates amyloid and TAU pathology in Alzheimer's brain. EMBO molecular medicine. 2016;8: 1005–1018. doi:10.15252/emmm.201606520

28. Che H, Sun L-H, Guo F, Niu H-F, Su X-L, Bao Y-N, et al. Expression of amyloid-associated miRNAs in both the forebrain cortex and hippocampus of middle-aged rat. Cellular Physiology and Biochemistry: International Journal of Experimental Cellular Physiology, Biochemistry, and Pharmacology. 2014;33: 11–22. doi:10.1159/000356646

29. Clovis YM, Enard W, Marinaro F, Huttner WB, De Pietri Tonelli D. Convergent repression of Foxp2 3'UTR by miR-9 and miR-132 in embryonic mouse neocortex: Implications for radial migration of neurons. Development (Cambridge, England). 2012;139: 3332–3342. doi:10.1242/dev.078063

30. Hinske LC, França GS, Torres HAM, Ohara DT, Lopes-Ramos CM, Heyn J, et al. miRIAD-integrating microRNA inter- and intragenic data. Database: The Journal of Biological Databases and Curation. 2014;2014. doi:10.1093/database/bau099

31. Chou C-H, Shrestha S, Yang C-D, Chang N-W, Lin Y-L, Liao K-W, et al. miRTarBase update 2018: A resource for experimentally validated microRNA-target interactions. Nucleic Acids Research. 2018;46: D296–D302. doi:10.1093/nar/gkx1067

32. Smalley SL, Loo SK, Yang MH, Cantor RM. Toward localizing genes underlying cerebral asymmetry and mental health. American Journal of Medical Genetics Part B, Neuropsychiatric Genetics: The Official Publication of the International Society of Psychiatric Genetics. 2005;135B: 79–84. doi:10.1002/ajmg.b.30141
