## Supplemental Table S5 for "Music-listening regulates human microRNA transcriptome"

| Comparison | Target genes of DE microRNAs that show overlap with the compared studies |
|---|---|
| Target genes of down-regulated microRNAs and singing stimulated genes from songbirds [1–3] | *ABCB7, ABCE1, ACTN1, ACTR3, ADAM22, ALG2, AMMECR1, ARL5B, ARPP19, ASF1A, ATP5G3, BBS10, BHLHE22, CRLF3, CTTNBP2, DCUN1D5, DGKH, DSCAM, DSTN, EGR3, EIF5A2, ERRFI1, ETNK1, FAM126B, FBXO28, FKBP5, FOSL2, FOXG1, FRMPD4, GCLM, GINS1, GOLT1B, GSPT1, H3F3B, HDHD2, HMGN3, HSPH1, IMPAD1, KCTD16, KIAA1107, KIAA1429, KLF2, KLHL8, LPHN3, LYSMD3, MLF1, MSTO1, MTAP, NUFIP2, NUP155, NUS1, OCIAD1, OXCT1, PAK1IP1, PCNX, PDCD10, PPIL3, PTS, RAB30, RAB6A, RABGEF1, RSL24D1, RSPO3, SAR1A, SCHIP1, SENP6, SFPQ, SGCZ, SLAIN2, SLC2A3, SLC30A1, SPOPL, STX7, SUB1, SUFU, TMEM106B, TMEM50B, TOB2, TRMT11, TTC33, TWSG1, UBE2B, UBE2D3, UFM1, USP16, VEGFA, WDR37, ZDHHC17, ZNF326, ZNF706* |
| Target genes of up-regulated microRNAs and singing inhibited genes from songbirds [2] | *AJAP1, ANK3, ANKRD17, ARHGEF9, ARID4B, ARL5B, ARPP19, ATP6V0A2, ATXN1, B3GALT2, BEND6, BTF3, CADM2, CAMK2D, CCDC88A, CDC42, CHST11, CISD1, CPSF4, CSMD1, DBT, DDX3X, DIRAS2, DNAJA2, DNAJB4, DNAJB5, DUSP6, EBAG9, EDEM3, EIF2S2, EIF3A, ELK4, EPC1, EYA1, FARP1, FGF14, FKBP1A, FUBP1, GRAMD1B, GRIK3, GRM7, GSPT1, HERC2, HIVEP2, HSP90AA1, HSPH1, INTS6, ISCA2, JHDM1D, JMY, KIAA2026, MED13L, MOCS2, MTF1, NAP1L1, NETO1, NLGN1, NUS1, PCLO, PDP2, PHF20L1, RGS7BP, RRP15, RWDD1, SAMD12, SCN2A, SERTAD4, SESTD1, SRPR, STXBP5, TAF1, TBL1XR1, TMEM120B, TNRC6B, TOP1, TRIOBP, TULP4, UACA, UNC13A, UQCRFS1, VGLL3, VPS35* |

### References

1. Wada K, Howard JT, McConnell P, Whitney O, Lints T, Rivas MV, et al. A molecular neuroethological approach for identifying and characterizing a cascade of behaviorally regulated genes. Proceedings of the National Academy of Sciences of the United States of America. 2006;103: 15212–15217. doi:10.1073/pnas.0607098103

2. Whitney O, Pfenning AR, Howard JT, Blatti CA, Liu F, Ward JM, et al. Core and region-enriched networks of behaviorally regulated genes and the singing genome. Science (New York, NY). 2014;346: 1256780. doi:10.1126/science.1256780

3. Hilliard AT, Miller JE, Fraley ER, Horvath S, White SA. Molecular microcircuitry underlies functional specification in a basal ganglia circuit dedicated to vocal learning. Neuron. 2012;73: 537–552. doi:10.1016/j.neuron.2012.01.005
